# Squash ensures Spindle-E–dependent heterotypic ping-pong amplification of piRNAs in the *Drosophila* ovary

**DOI:** 10.64898/2025.12.12.693873

**Authors:** Xin Xu, Taichiro Iki, Shinichi Kawaguchi, Toshie Kai

## Abstract

The PIWI-interacting RNA (piRNA) pathway plays a crucial role in repressing mobile transposable elements (TEs) and protecting the integrity of the heritable genome in animal gonads. In the *Drosophila* ovary, piRNAs are produced in a membrane-less organelle called nuage, in which many piRNA factors are localized. Among them, Squash (Squ) has also been identified as a key component in piRNA-directed TE silencing. However, its molecular function remains largely unknown. Here, we demonstrate that loss of Squ leads to defective piRNA biogenesis, which is associated with the abnormal accumulation of precursor transcripts and the specific destabilization of Ago3, a member of the PIWI-family proteins. Reducing Ago3 results in enhanced homotypic piRNA amplification mediated by Aub itself, rather than the heterotypic ping-pong amplification between Aub and Ago3. Additionally, we demonstrate that Squ is a functional cofactor of the RNA helicase, Spindle-E (Spn-E), which plays a vital role in piRNA biogenesis. Point mutations that abrogate the interaction between Squ and Spn-E result in TE de-repression and defective precursor processing, suggesting that Squ works with Spn-E to ensure the proper piRNA biogenesis. These results identify Squ as a critical factor for Spn-E–dependent heterotypic ping-pong amplification of piRNAs in the *Drosophila* ovary.

## Introduction

Transposable elements (TEs) are mobile genetic elements that can replicate and insert into new genomic locations, present in most eukaryotic organisms (Wells and Feschotte, 2020). Although TEs have contributed substantially to the evolution of genomes, their uncontrolled mobilization poses a major threat to genome integrity by disrupting gene expression and inducing DNA damage (Kazazian, 2004; Goodier and Kazazian, 2008; Hedges and Belancio, 2011; Betancourt et al., 2024). In the gonads of *Drosophila melanogaster*, TE activity is repressed by PIWI-interacting RNAs (piRNAs), a class of 23–29 nt small non-coding RNAs that associate with PIWI-clade Argonaute (Ago) family proteins, Piwi, Aubergine (Aub), and Ago3 to form piRNA-induced silencing complexes (piRISCs). These complexes mediate both transcriptional and post-transcriptional silencing of TEs, thereby safeguarding germline genome integrity (Brennecke et al., 2007; Czech and Hannon, 2011; Darricarrère et al., 2013).

piRNAs are derived from long precursor transcripts generated at discrete genomic loci known as piRNA clusters, which serve as reservoirs of TE fragments and subsequently produce antisense piRNAs against active elements (Brennecke et al., 2007; Malone and Hannon, 2009). In *Drosophila*, piRNA clusters are classified into two distinct clusters: dual-strand clusters which are transcribed from both genomic strands, and uni-strand clusters, transcribed from a single strand. The *flamenco* locus functions as the major uni-strand cluster in ovarian somatic cells, whereas *42AB* and *38C* represent the predominant germline dual-strand clusters that contribute to most piRNA production in ovarian germline cells (Brennecke et al., 2007; Goriaux et al., 2014; Andersen et al., 2017). In *Drosophila* germline cells, these dual-strand piRNA precursors are exported from the nucleus to specific membrane-less granules termed nuage in the perinuclear cytoplasmic region, mediated by the Nxf3-Nxt1 export pathway (ElMaghraby et al., 2019; Kneuss et al., 2019; Mendel and Pillai, 2019; Lin et al., 2023). piRNA precursors and transposon transcripts are subsequently loaded onto Aub and Ago3 by the function of RNA helicases such as Vas to initiate the amplification process known as the ping-pong cycle (Xiol et al., 2014). During this process, Aub and Ago3 engage in reciprocal cleavage of target transcripts with a 10-nt overlap at 5’-ends, producing sense and antisense piRNAs for repressing TEs (Brennecke et al., 2007; Gunawardane et al., 2007; Aravin et al., 2007; Lim et al., 2014).

In addition to Aub and Ago3, a set of nuage-localizing proteins are indispensable for precursor processing and ping-pong amplification of piRNAs. Notably, Tejas (Tej), a Tudor-domain-containing protein, functions as a molecular scaffold that recruits the RNA helicases Vasa (Vas) and Spindle-E (Spn-E) to nuage, thereby facilitating both piRNA precursor processing and the dynamic assembly of functional piRNA complexes (Lim and Kai, 2007; Nishida et al., 2007; Malone et al., 2009; Patil and Kai, 2010; Xiol et al., 2014; Sato et al., 2015; Lin et al., 2023; Adashev et al., 2024). Other nuage components, such as Krimper (Krimp) and Qin/Kumo, also act as regulators of PIWI-family protein interactions and strand-biased production of piRNAs during ping-pong amplification (Zhang et al., 2011; Anand and Kai, 2012; Sato et al., 2015). Taken together, these proteins form a highly coordinated network within the nuage that ensures robust piRNA biogenesis.

By contrast, the molecular functions of certain nuage components remain poorly understood. Squash (Squ), a *Drosophila*-specific protein, was initially proposed as an RNAse HII-related protein (Pane et al., 2007), yet structural prediction by the AlphaFold program suggested that it is an intrinsically disordered protein lacking any conserved or characterized motifs/domains (Kawaguchi et al., 2025). Squ has been implicated to function in piRNA biogenesis in *Drosophila*: upon loss of *squ* function, upregulation of some TEs and a mild reduction of total piRNAs are observed, while no severe disruption for the localization of core piRNA components is detected (Pane et al., 2007; Malone et al., 2009; Kawaguchi et al., 2025). Although *squ* mutant females can produce eggs, a range of dorsoventral (DV) patterning defects are observed in mutant oocytes and the eggs from *squ* mutant mother cannot hatch (Schüpbach and Wieschaus, 1991; Pane et al., 2007). In this study, we identified Squ as a crucial partner of Spn-E, orchestrating the ping-pong amplification of piRNAs and piRNA processing from Aub to Ago3 at nuage. Also, Squ sustains controlled heterotypic ping-pong amplification between Aub and Ago3 by maintaining Ago3 protein level in *Drosophila* germline cells.

## Results

### Nuage component Squ is specifically required for Ago3 abundance

Previous studies reported that loss of *squ* reduces total piRNAs by ∼50% without markedly altering the nuage localization of core ping-pong factors such as Aub and Ago3 (Pane et al., 2007; Malone et al., 2009). Immunostaining of the control ovaries showed that endogenous Squ was enriched in the perinuclear nuage granules of nurse cells (Fig. 1A). This finding was consistent with previous observations using a transgene expressing HA-tagged Squ, confirming the localization of Squ to the nuage (Pane et al., 2007). To further examine the molecular role of Squ, we analyzed the localization and abundance of major piRNA pathway components in *squ* mutant ovaries using the trans-heterozygous alleles, *squ*^PP32/HE4*7*^(hereafter, *squ* mutant). In *squ* mutant ovaries, the perinuclear signals of most core ping-pong factors (Spn-E, Aub, Vas and Tej) were largely unaffected (Fig. 1A). Reversely, in the mutant of *spn-E*, *vas*, or *tej*, Squ protein level was decreased and detected only faintly in the cytoplasm (Fig. 1B, C and Fig. S1A, B), indicating that Squ is not an indispensable factor for nuage formation but rather a factor recruited by these core ping-pong factors to nuage.

**Figure 1.**
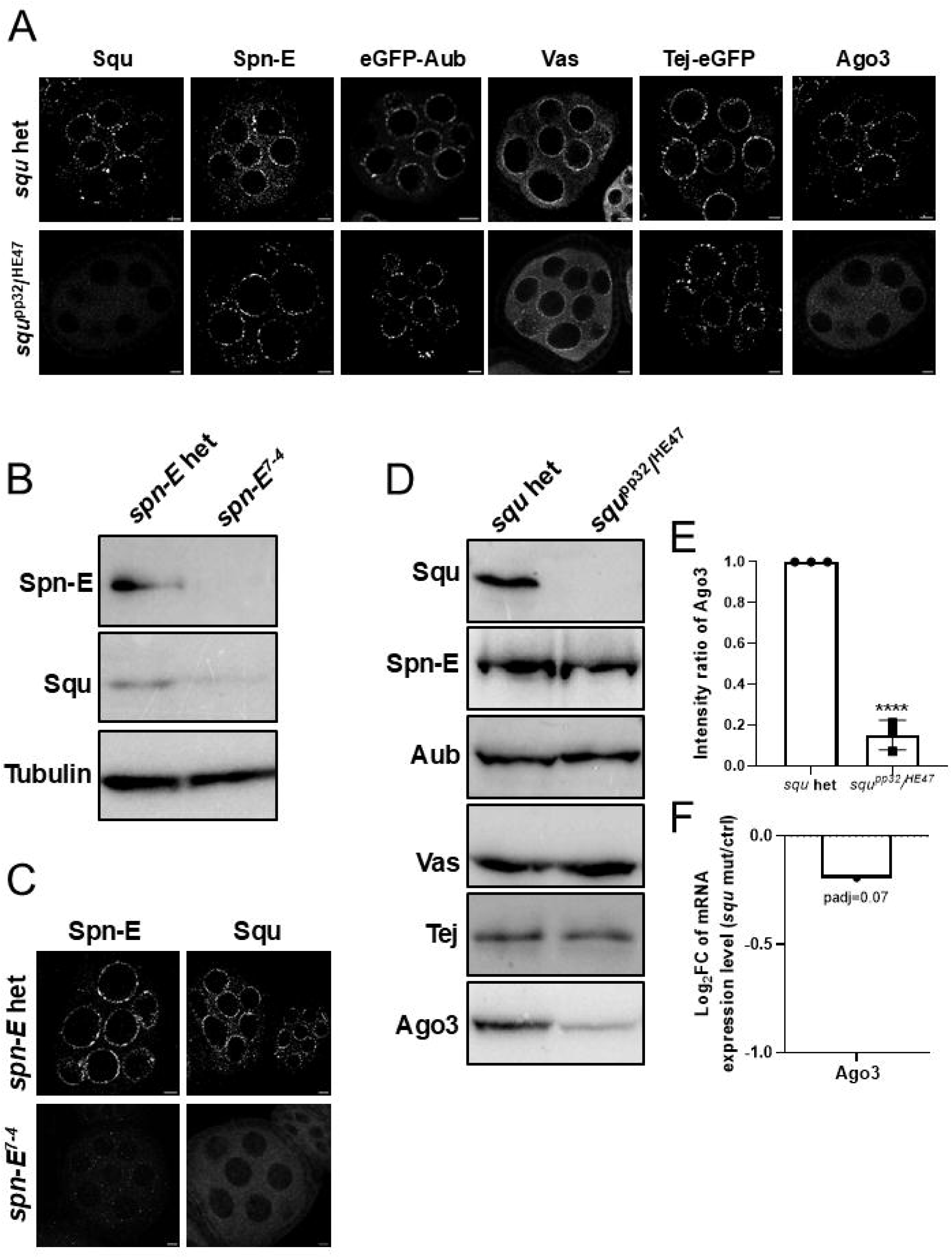
Squ is required for Ago3 abundance and exhibits *spn-E*–dependent nuage enrichment. (A) Localization of indicated proteins in stage 4-5 egg chambers of *squ* heterozygous or transheterozygous mutants. Immunostaining; Squ, Spn-E, Vas, Ago3. Fluorescence; eGFP-Aub and Tej-eGFP. Scale bar is 5 μm. (B) Protein level of Squ in control and *spn-E* mutant egg chambers. (C) Immunostaining of Squ in control and *spn-E* mutant egg chambers. Scale bar is 5 μm. (D) Immunoblotting of indicated proteins extracted from ovaries of *squ* heterozygous or mutants. (E) Quantification of Ago3 protein signal intensities. Error bars indicate standard deviation (3 biological replicates). Statistics: unpaired t-test, ****p < 0.0001. (F) Ago3 transcript levels in control and *squ* mutant ovaries. Fold change given by differential expression analysis (DESeq2) is shown with adjusted p-value.

We found, in the immunofluorescence observations of *squ* mutant ovaries, Ago3 signal is markedly weak (Fig. 1A). Consistently, immunoblotting of *squ* mutant ovarian lysate revealed a reduction in Ago3 protein level by ∼80% compared to those in wild-type condition, whereas Aub, Spn-E, Vas and Tej were largely unchanged (Fig. 1D and E). Hence, this reduction is Ago3-specific. Ovarian transcriptome analysis confirmed that *ago3* mRNA level was essentially unchanged in *squ* mutant ovaries (log_2_FC = -0.19, padj = 0.07; Fig. 1F). Taken together, these data suggest that *squ* is critical in maintaining Ago3 protein abundance in germline cells.

### *squ* loss derepresses germline-specific transposons

Given the severe loss of Ago3 in *squ* mutant ovaries (Fig. 1), TEs might be derepressed. Indeed, RNA-seq analysis revealed that many TEs were significantly up-regulated in *squ* mutant ovaries (24 out of 119 TE families showed log₂FC > 1, p-value < 0.05; Fig. 2A). Consistent to the expression of Squ in germline cells (Fig. 1A), these upregulated TEs mostly belonged to germline-specific families (Fig. 2B, S2A; Durdevic et al., 2018; Kneuss et al., 2019). RT-qPCR confirmed significant up-regulation of representative germline TEs, *HMS-Beagle*, *Burdock* and *blood*, in *squ* mutant ovaries (Fig. 2C). As an independent protein-level readout of de-repression, we examined a canonical piRNA-controlled target, HeT-A, by immunostaining and detected robust increase of HeT-A Gag protein in oocyte of *squ* mutant ovaries—which is normally silenced by the piRNA pathway (Fig. 2D; Shpiz et al., 2011). Together, these data indicate that *squ* is required to repress a broad set of germline TEs.

**Figure 2:**
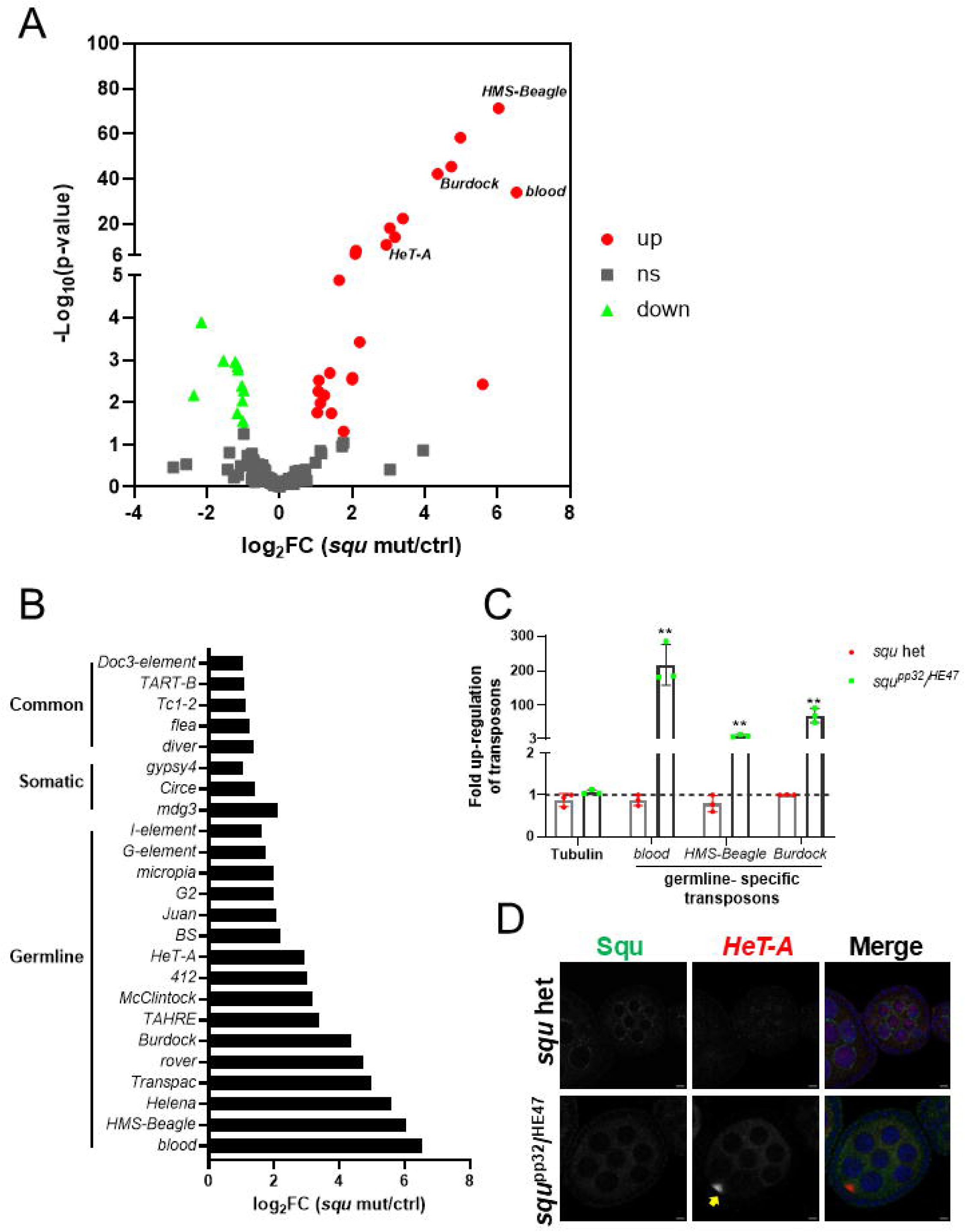
Loss of *squ* derepresses transposons. (A) Volcano plot shows fold changes of transposon transcript levels in ovaries (*squ* mutant compared to *squ* heterozygous condition). Red circles; significantly upregulated transposons (log_2_FC > 1, p < 0.05), greed circles; significantly downregulated transposons (log_2_FC < −1, p < 0.05). (B) Fold changes of TEs’ expression upon *squ* loss. Transposons are categorized into germline, somatic, and common groups. (C) RT-qPCR analysis of germline-specific transposons (*blood*, *HMS-Beagle*, *Burdock*). Relative transcript levels are calculated by ^ΔΔ^Ct using rp49 as reference. Values represent fold changes in *squ* mutant relative to heterozygous controls. Error bars indicate standard deviation (3 biological replicates). Statistical analysis is performed using Student’s t-test. **p < 0.01. (D) Immunostaining of HeT-A Gag and Squ proteins in control and *squ* mutant ovaries. A yellow arrow indicates HeT-A Gag signals accumulating in oocyte. Scale bar: 5 μm.

### Squ is required for maintenance of Ago3-bound piRNAs and stalls piRNA precursor processing at nuage

The de-repression of TEs always represents the reduction of piRNA production and previous study has already reported that the total number of piRNAs was reduced to half in *squ* mutant ovaries (Malone et al., 2009). However, the cause of the reduction in piRNAs remain unclear. To address this question, we analyzed 23-29 nt small RNAs bound to Aub and Ago3. NGS analysis revealed that the amount of Ago3-bound piRNAs from piRNA clusters *38C-3* and *42AB* were strongly depleted in *squ* mutant ovaries but that of Aub-bound piRNAs were not largely affected (Fig. 3A, rep1; Fig. S3A, rep2). Although without significant reduction for Aub-bound piRNAs, the antisense bias of Aub-bound piRNAs mapping to transposons was reduced from ∼77% to ∼55% in *squ* mutant ovaries (Fig. 3B and S3B). Of those, Aub-bound piRNAs mapping to severely upregulated TEs in *squ* mutant ovaries, such as *HMS-Beagle*, *blood*, *Burdock* and *HeT-A*, showed severe reduction of antisense bias (Fig. 3C and S3C), implying that this reduction of antisense bias may further result in upregulation of corresponding TEs. Take together, these data suggest that Squ is required for efficient production for Ago3-bound piRNAs, which in turn maintains the antisense enrichment of Aub-bound piRNAs for effective transposon silencing.

**Figure 3:**
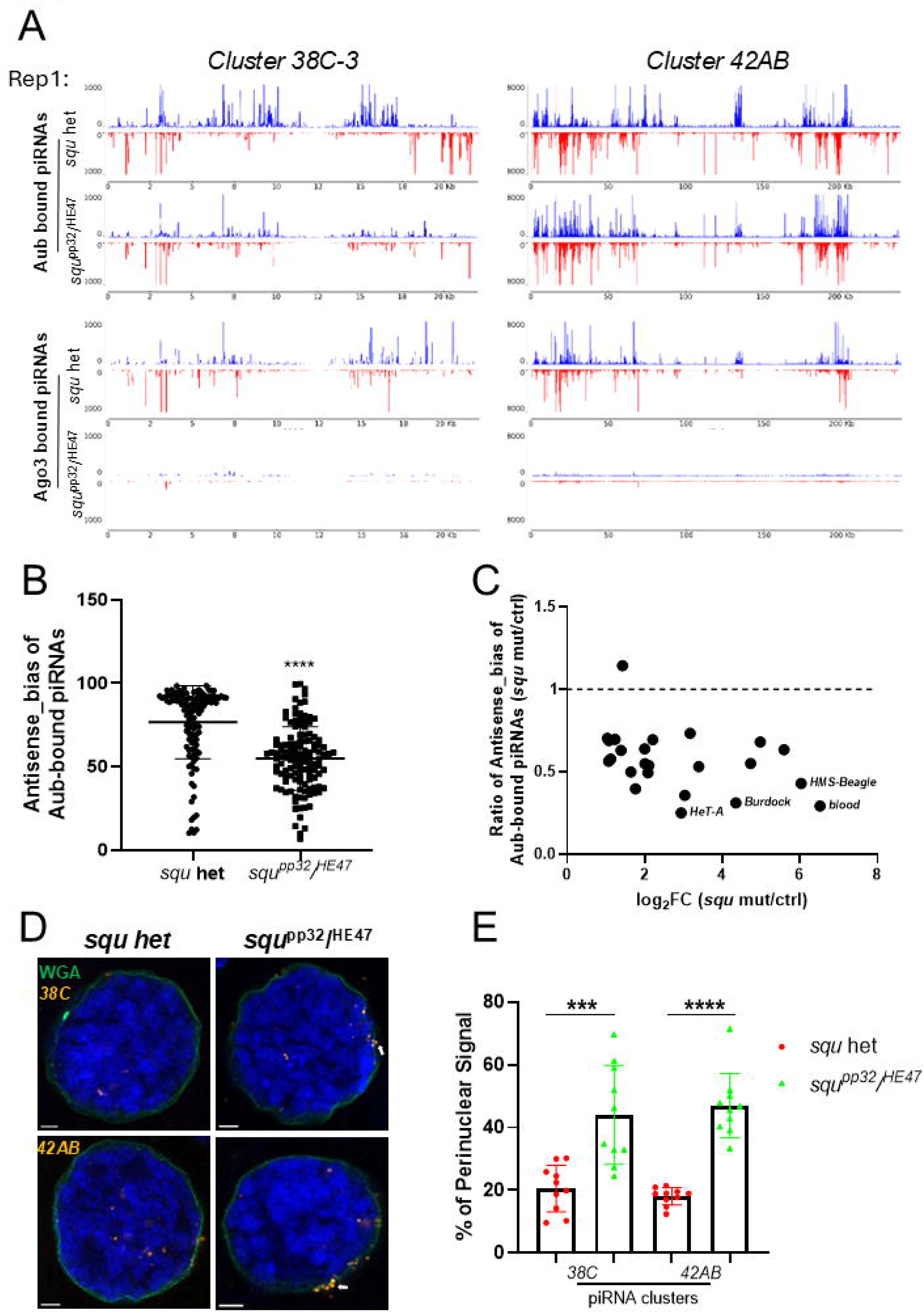
Squ is required for maintenance of Ago3-bound piRNAs and processing of piRNA precursors. (A) Aub- (upper panels) and Ago3-bound (lower panels) small RNAs from control and *squ* mutant ovaries mapped to the major piRNA clusters *38C-3* and *42AB* (replicate 1). Plus-strand (blue) and minus-strand (red) piRNAs are shown as upward and downward peaks, respectively. (B) Antisense bias of Aub-bound piRNAs mapping to transposable elements (TEs) in control (*squ* het) and *squ* mutant ovaries (replicate 1). Each dot represents one TE; antisense piRNA ratios are calculated by dividing antisense-mapping piRNA reads by total mapping reads for each TE. Horizontal lines indicate the mean values. ****p < 0.0001 (unpaired t-test). (C) Effect of loss of *squ* function on antisense ratio of Aub-bound piRNAs mapping to transposon consensus sequences. Dots indicate 24 up-regulated TEs (Fig. 2A) in *squ* mutant relative to control ovaries (replicate 1). For each TE, the antisense bias in *squ* mutant is divided by that in *squ* heterozygous. (D) HCR-FISH detection of piRNA precursor transcripts from clusters *38C* and *42AB* in control and *squ* mutant ovaries. One representative nucleus for each sample is shown. A white arrow indicates precursor signals accumulating to perinuclear regions. Blue; DNA (DAPI), green; nuclear membrane (WGA). Scale bar: 2 μm. (F) Quantification of piRNA precursor signals accumulating in perinuclear regions. Fluorescence intensity of precursor foci within ±5% of nuclear diameter from the nuclear membrane is measured (n=10 nuclei per genotype). Signals are plotted as the percentage of total nuclear-associated intensity. Statistical analysis: *** p < 0.001, **** p < 0.0001.

As precursor processing disruption always results in piRNA reduction, we next examined whether the recruitment of piRNA precursors to perinuclear nuage was perturbed upon loss of *squ* function. Hybridization chain reaction–fluorescence in situ hybridization (HCR–FISH; Choi et al., 2018) targeting precursors from the major dual-strand clusters in germline cells, *38C* and *42AB*, showed accumulated perinuclear signals in *squ* mutant nurse cells compared with controls (Fig. 3D). Quantification using a perinuclear-rim index (Lin et al., 2023) revealed that ∼45% of the total precursor signal resided at the perinuclear rim in *squ* mutant nurse cells vs ∼20% in heterozygous controls (Fig. 3E). This perinuclear accumulation in *squ* mutant ovaries is comparable to that reported for *vas*, *tej*, and *spn-E* mutants, which function on recruiting precursors or TE transcripts to initiate the ping-pong amplification at nuage (Lin et al., 2023). Consistently, piRNA-cluster transcripts were increased in *squ* mutant ovaries by RT-qPCR (Fig. S3D). Because signals remained predominantly cytoplasmic rather than nuclear, these observations argue against a primary nuclear-export defect and instead suggest inefficient precursor processing in perinuclear nuage upon loss of *squ* function. Taken together, these findings place *squ* as a factor required for Ago3-bound piRNA production and the efficient perinuclear processing of piRNA precursors.

### Low Ago3 protein level in *squ* mutant triggers Aub-Aub homotypic ping-pong

In *Drosophila* germline cells, the canonical ‘ping-pong’ amplification loop also known as ‘heterotypic ping-pong’ occurs between Aub (antisense, 1U bias for bound piRNA) and Ago3 (sense, 10A bias for bound piRNA), but in *kumo*/*qin* and *krimp* mutants, different extents of reduction for Ago3 protein level were detected together with a piRNA production step called ‘homotypic ping-pong’ occurred by Aub with itself harboring a ‘1U10A’ bias for bound piRNAs (Zhang et al., 2011; Sato et al., 2015). To test whether the low Ago3 level elicits Aub-Aub homotypic ping-pong, we examined piRNA profiles in *squ* and *ago3* mutant ovaries. The 1U nucleotide bias of Aub-bound piRNAs and 10A bias of Ago3-bound piRNAs were detected in controls, as expected (Fig. 4A). In contrast, Aub-bound piRNAs in *squ* mutant ovaries exhibited a strong ‘1U10A’ signature, a hallmark of Aub-Aub homotypic ping-pong (Fig. 4A and S4A), whereas Ago3-bound reads were too few to assess. Importantly, in *ago3* mutant ovaries, we also observed a strong ‘1U10A’ signature for Aub-bound piRNAs (Fig. 4B and S4B), the characteristic for Aub-Aub homotypic ping-ping, together with the reduction for antisense bias in line with the previously reported up-regulation of transposons like *HMS-Beagle*, *blood*, *Burdock* and *HeT-A* (Fig. 4C-E and S4C-E; Li et al., 2009). These data suggest that the role of Squ in maintaining Ago3 protein level is crucial for sustaining the heterotypic ping-pong cycle in *Drosophila* germline cells.

**Figure 4:**
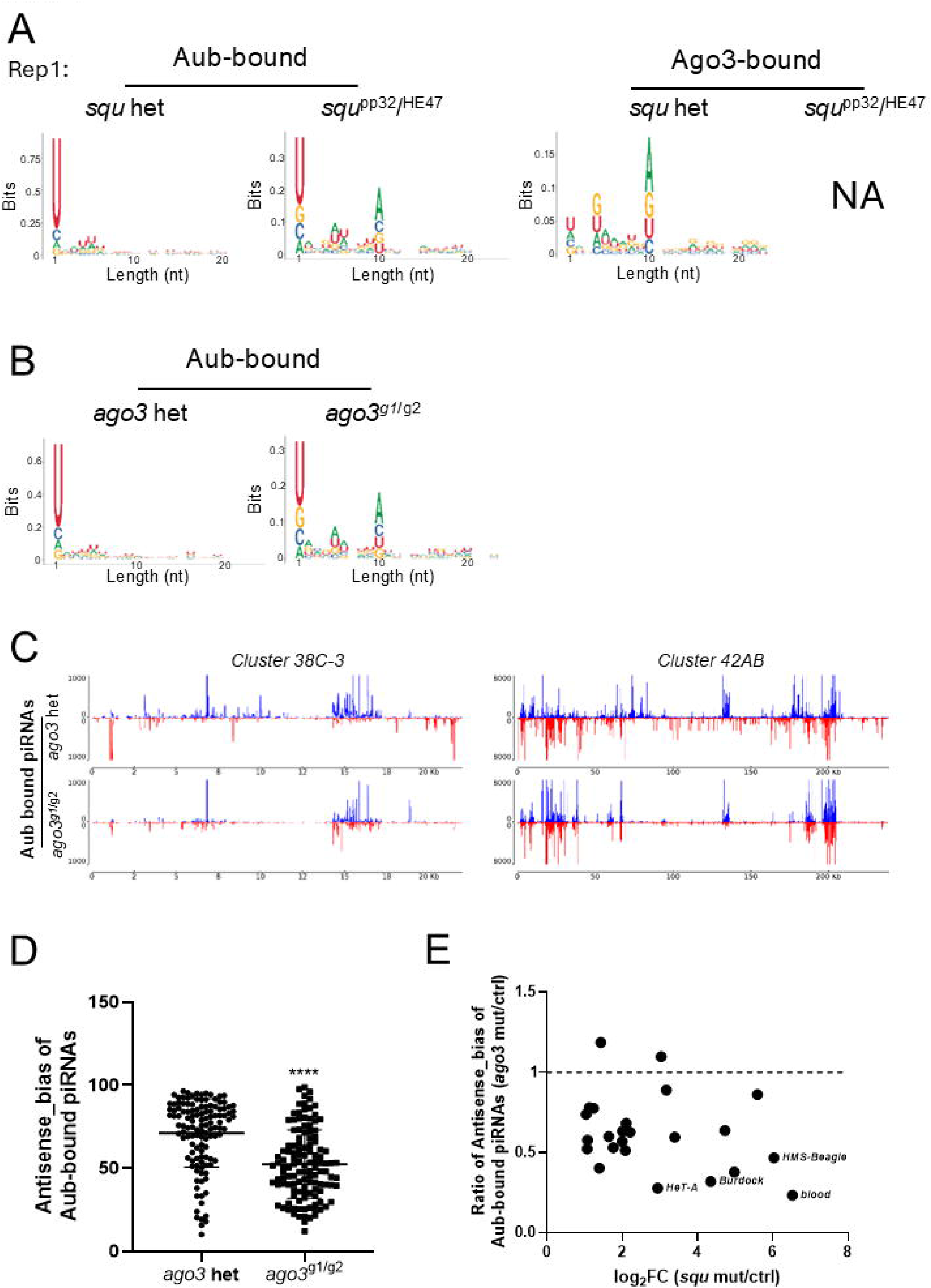
Low Ago3 protein level in *squ* mutant triggers Aub–Aub homotypic ping-pong. (A) Nucleotide bias of Aub-bound and Ago3-bound piRNAs mapped to piRNA cluster loci in control and *squ* mutant ovaries (replicate 1). Sequence logos indicate 1^st^ to 20^th^ of piRNAs. 10A signature of *squ* mutant condition reflects enhanced Aub–Aub homotypic ping-pong. NA (Not analyzed); Ago3-bound piRNAs are not detectable in *squ* mutant. (B) Nucleotide bias of Aub-bound piRNAs in control and *ago3* mutant ovaries (replicate 1). (C) Aub-bound piRNAs from control and *ago3* mutant ovaries mapped to the major germline piRNA clusters *38C-3* and *42AB*. Blue; genomic plus-strand reads, red; minus-strand reads. (D) Antisense bias of Aub-bound piRNAs mapping to transposable elements (TEs) in control and *ago3* mutant ovaries (replicate 1). Each dot represents one TE; antisense piRNA ratios are calculated by dividing antisense-mapping piRNA reads by total mapping reads for each TE. Horizontal lines indicate the mean values. ****p < 0.0001 (unpaired t-test). (E) Ratio of antisense bias of Aub-bound piRNAs for the 24 up-regulated TEs (detected in *squ* mutant) in *ago3* mutant versus control ovaries (replicate 1). Dots indicate 24 up-regulated TEs (Fig. 2A) in *ago3* mutant relative to control ovaries (replicate 1). For each TE, the antisense bias in *ago3* mutant is divided by that in *ago3* heterozygous.

### Squ–Spn-E interaction is essential for piRNA precursor processing and TE repression

In previous studies, Squ protein was detected, together with several piRNA factors, in immunoprecipitates of Spn-E from ovarian lysates, indicating that Squ and Spn-E interact either directly or indirectly (Andress et al., 2016). Recently, AlphaFold2 predicted their direct interaction with high confidence, and co-immunoprecipitation assays in S2 culture cells confirmed this specific interaction between Squ and Spn-E (Kawaguchi et al., 2025). Consistently, Squ co-immunoprecipitated with Spn-E from ovarian lysates, reflecting a specific interaction *in vivo* (Fig. 5A). A faint signal with Ago3 was also observed (Fig. 5A and S5A), whereas Aub, Vas, and Tej were undetectable in the Squ-immunoprecipitants, indicating that Squ preferentially associates with Spn-E, and only a minor potential interaction with Ago3.

**Figure 5:**
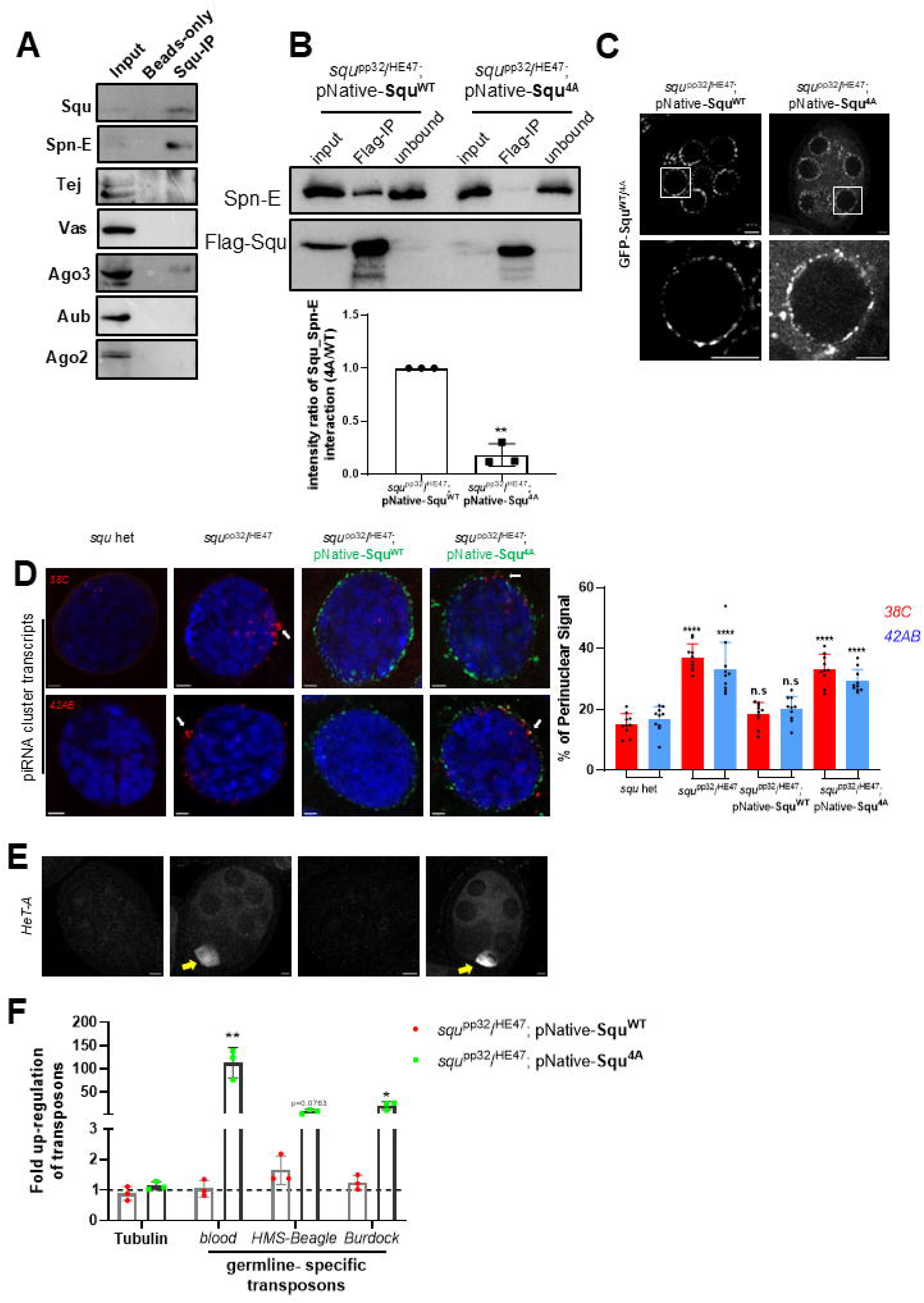
Interaction between Squ and Spn-E is crucial for piRNA biogenesis, transposon silencing, and oogenesis. (A) Immunoblotting analysis of Squ interactors in ovaries. Squ is immunoprecipitated from the control *y w* ovaries using anti-Squ antibodies. Precipitation process without anti-Squ antibodies serves as negative control. piRNA biogenesis factors; Spn-E, Tej, Vas, Ago3, and Aub. Ago2, not involved in piRNA biogenesis, serves as a negative control. (B) Immunoprecipitation of GFP–3×Flag–Squ and the variants expressed under the native promoter (pNative-Squ^WT^ and pNative-Squ^4A^) using anti-3FLAG antibodies. The lower panel shows the quantification of the Squ–Spn-E interaction, plotted as the intensity ratio relative to Squ^WT^ (3 biological replicates). (C) Localization of Squ^WT^ and Squ^4A^ expressed under the native promoter in *squ* mutant germline cells (upper panels: stage 4–5 egg chambers; lower panels: magnified nuclei indicated by squares in the upper panels). Squ^WT^ is enriched in perinuclear nuage, whereas Squ^4A^ shows detached aggregates in the cytoplasm in addition to the perinuclear foci. Scale bar, 5 μm. (D) HCR in situ hybridization of the major germline piRNA precursors derived from clusters *38C* and *42AB* in germline cells of heterozygous control, *squ*^PP32/HE47^, and *squ*^PP32/HE47^ expressing Squ^WT^ or Squ^4A^ under the native promoter. DNA is stained with DAPI (blue); the nuclear membrane is labeled with WGA (green). White arrows indicate precursor signals accumulating to perinuclear regions. Scale bar, 2 μm. Right, quantification of perinuclear precursor signals. Fluorescence intensities of foci located within ±5% of the nuclear diameter inside and outside the nuclear membrane are summed, normalized to the total nuclear signal, and plotted as the percentage of perinuclear signal. (E) Immunostaining of HeT-A Gag protein in the heterozygous control, *squ*^PP32/HE47^mutant, and *squ*^PP32/HE47^ expressing Squ^WT^ or Squ^4A^ ovaries. Yellow arrows indicate HeT-A Gag signals. Scale bar, 5 μm. (F) RT–qPCR analysis of transcripts derived from the germline-specific transposons *blood*, *HMS-Beagle*, and *Burdock*, with tubulin as a control. All values are normalized to *rp49* and are shown relative to expression levels in *squ*^PP32/HE47^; pNative-Squ^WT^ ovaries. Error bars indicate standard deviation (3 biological replicates). Statistical significance is assessed by Student’s unpaired t-test. *p < 0.05, **p < 0.01.

To investigate the molecular basis and physiological relevance of the Squ–Spn-E interaction, we first examined the interface predicted by AlphaFold2. Four residues in Squ (E107, E109, R115, and K163), which are conserved in *Drosophila* Squ homologs shown in our previous study (Kawaguchi et al., 2025), are predicted to form salt bridges with four corresponding residues in Spn-E (K834, R675, D640, and E1274). Substituting these four residues with alanine in Squ (Squ^4A^) abolished its interaction with Spn-E in S2 cells (Kawaguchi et al., 2025). To complement this, in this study, we generated Spn-E mutant with alanine substitutions at the predicted interface residues (Spn-E^4A^), which likewise lost binding to Squ in co-immunoprecipitation assays in S2 culture cells (Fig. S5B), collectively confirming that these conserved residues mediate the interaction via salt bridges.

To assess the physiological consequences of disrupting this interaction *in vivo*, Squ^4A^ or wild-type Squ (Squ^WT^) was expressed under the native promoter in ovaries lacking endogenous *squ*. Co-immunoprecipitation revealed markedly reduced binding of Squ^4A^ to Spn-E compared with Squ^WT^ in ovaries (Fig. 5B). Besides, Squ^4A^ showed detached aggregates in the cytoplasm in addition to the perinuclear foci and was expressed at lower levels than Squ^WT^ (Fig. 5C and S5C). Rescue experiments further demonstrated that Squ^WT^ restored proper localization of piRNA precursors, processing of piRNA cluster transcripts as reflected by reduction of their levels (Fig. 5D and S5D), repression of TEs and suppression of HeT-A Gag protein accumulation (Fig. 5E and F). Moreover, Squ^WT^ restored female fertility as measured by hatching rates (Fig. S5E). In contrast, Squ^4A^ failed to rescue any of these defects despite with a higher expression than endogenous Squ (Fig. 5C-F and S5C-E). Collectively, these results indicate that the four conserved residues in Squ are critical for direct binding to Spn-E *in vivo* and this interaction is essential for proper piRNA biogenesis, TEs’ repression, and female fertility in *Drosophila*.

### Squ contributes to Ago3 protein maintenance independently of Spn-E

In *squ* mutant ovaries, both Ago3 immunostaining signals and total protein levels were severely reduced (Fig. 1). This reduction was efficiently rescued by expression of Squ^WT^, whereas the Squ^4A^ mutant protein restored Ago3 only weakly (Fig. 6A and B), suggesting that Squ contributes to the maintenance of Ago3 protein levels. To further examine whether the Squ–Spn-E complex is required for the Ago3 stability or Squ has a direct effect on maintaining Ago3 protein, Ago3 localization and protein levels were analyzed by expressing Squ^WT^ or Squ^4A^ in the *spn-E* mutant background. Previous studies have reported a strong reduction of Ago3 protein in *spn-E* mutant ovaries (Ryazansky et al., 2016; Andress et al., 2016), which we also confirmed with our newly generated loss of function allele, *spn-E^7-4^*, in this study. Upon the loss of *spn-E* function, Ago3 protein level was strongly reduced together with a pronounced decrease in endogenous Squ (Fig. 6C). Notably, the expression of Squ^WT^ substantially increased Ago3 abundance in *spn-E* mutant ovaries, suggesting that Squ^WT^ stabilizes Ago3 protein in a Spn-E-independent manner. On the other hand, the Squ^4A^ mutant had only a minor effect on the level of Ago3, which may be due to its lower expression level and/or the impaired functional activity of the mutant protein (Fig. 6C).

**Figure 6:**
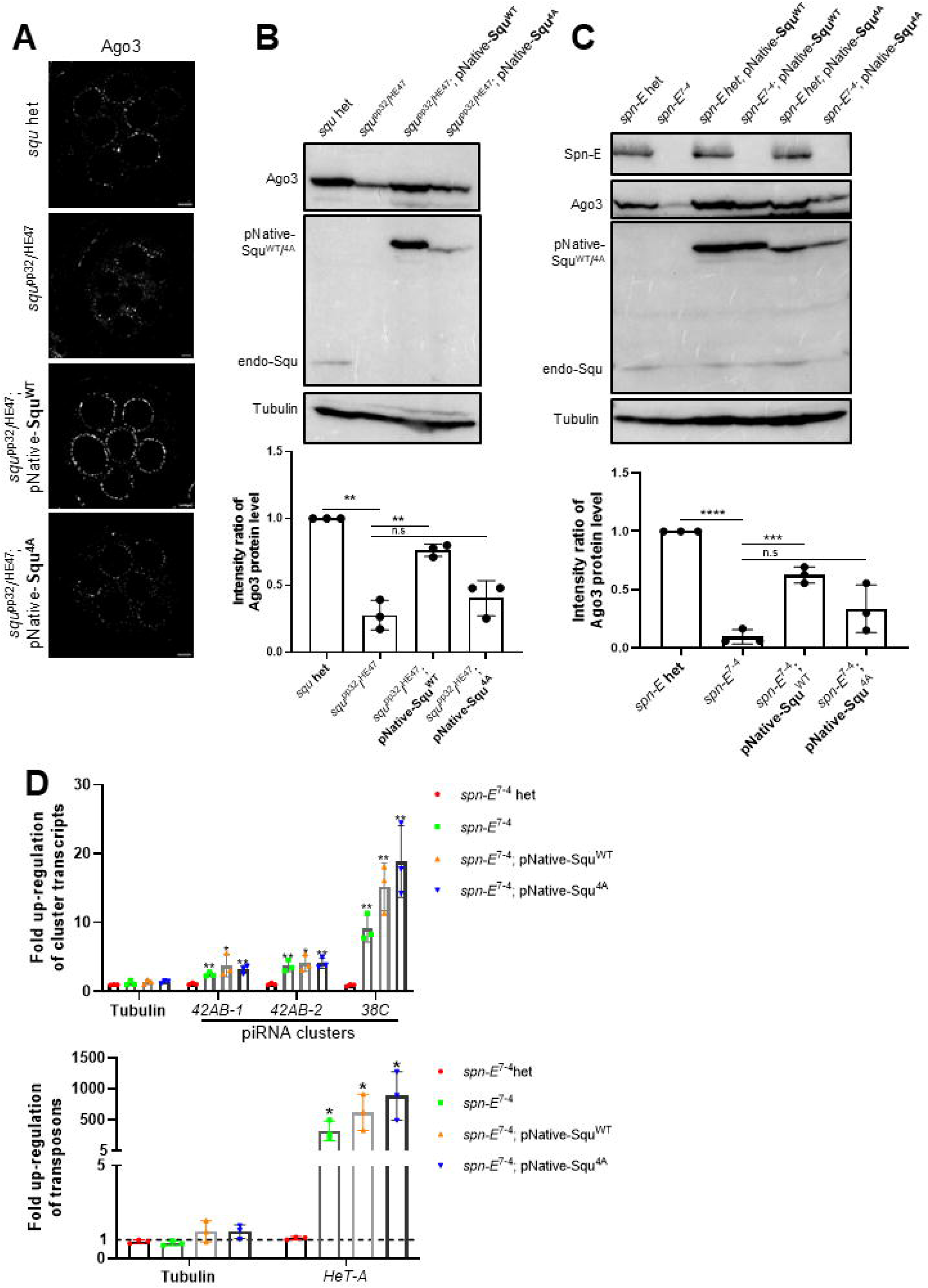
Squ can stabilize Ago3 without Spn-E. (A) Immunostaining of mid-stage egg chambers from the heterozygous control, *squ*^PP32/HE47^, and *squ*^PP32/HE47^ expressing Squ^WT^ or Squ^4A^ under the native promoter. Scale bar is 5 μm. (B) Immunoblotting of proteins extracted from the heterozygous control, *squ*^PP32/HE47^ and *squ*^PP32/HE47^expressing Squ^WT^ or Squ^4A^ under the native promoter. Lower panel shows the quantification of signal intensities (3 biological replicates). Statistical significance is assessed by Student’s unpaired t-test. **p < 0.01. (C) Immunoblotting of proteins extracted from the ovaries of the heterozygous control, *spn-E*^7-4^, and *spn-E*^7-4^ expressing Squ^WT^ or Squ^4A^ under the native promoter. Lower panel shows the quantification of signal intensities (3 biological replicates). Statistical significance is assessed by Student’s unpaired t-test. ***p < 0.001, ****p < 0.0001. (D) RT-qPCR measurement of transcripts from piRNA cluster loci, *38C* and *42AB*, those from *HeT-A* and the control tubulin. Relative expression values are obtained by ^ΔΔ^Ct using *rp49* as reference. Error bars; standard deviation of biological triplicates. Statistics; unpaired T-test (*p <0.05, **p <0.01).

To test whether Squ associates with Ago3 independently of Spn-E, we performed co-immunoprecipitation assays in the *spn-E* mutant background. Both Squ^WT^ and Squ^4A^ were found to associate with Ago3 even in the absence of Spn-E (Fig. S6A), indicating that Squ can interact with Ago3 independently of Spn-E. In contrast, overexpression of Flag-Ago3 by a strong germline driver, NGT40 and nosGal4, in *squ* mutant ovaries failed to restore Ago3 protein levels (Fig. S6B), indicating that Squ is required for Ago3 protein stabilization rather than for Ago3 expression itself. Together, these results suggest that although proper Squ–Spn-E function is required for robust maintenance of Ago3 protein, Squ also exerts a Spn-E–independent effect on Ago3 protein stability through an as-yet-unknown mechanism.

To determine whether the defects in piRNA precursor processing and transposon repression in *spn-E* mutant ovaries are primarily caused by Ago3 reduction, we analyzed the expression of piRNA cluster transcripts from *38C* and *42AB*, as well as the retrotransposon *HeT-A* in *spn-E* mutant ovaries (Fig. 6D). Despite the substantial restoration of Ago3 protein levels by Squ^WT^ expression, the upregulation of cluster transcripts and *HeT-A* remained largely unchanged in *spn-E* mutant ovaries (Fig. 6D). Similar results were observed with Squ^4A^ (Fig. 6D). These results indicate that recovery of Ago3 protein alone is not sufficient to rescue the defects in piRNA precursor processing or transposon repression in *spn-E* mutant ovaries. Instead, it implies that Spn-E acts upstream of both Squ and Ago3 in the piRNA pathway, supporting the notion that Spn-E is a core nuage factor essential for piRNA pathway activity.

### Squ facilitates Spn-E–mediated RNA processing at Aub

A recent study in silkworm demonstrated that Spn-E is recruited to the Siwi (the silkworm ortholog of *Drosophila* Aub)–piRNA–target RNA complex formed with GTSF1 and Meal (referred to as Siwi* or Aub*) (De et al., 2025). To examine whether the formation or dynamics of the Aub*–Spn-E complex is affected by Squ, we analyzed the interaction between Aub and Spn-E by immunoprecipitation. When Aub was immunoprecipitated from control ovary lysates, Spn-E was barely detectable in the bound fraction. In contrast, the amount of Spn-E co-precipitated with Aub was increased in *squ* mutant ovaries (Fig. 7A and S7A). These results indicate that Spn-E remains more stably associated with Aub* in the absence of Squ. To further examine the nature of this Aub*–Spn-E complex, we analyzed the RNA species recovered in the Spn-E immunoprecipitants. The levels of piRNA cluster transcripts derived from *38C* and *42AB*, which serve as major target RNAs for Aub, were significantly increased in *squ* mutant ovaries compared with the heterozygous controls, despite a similar amount of Spn-E being present in the IP fractions (Fig. 7B and S7B). These data indicate that, in the absence of Squ, Spn-E is retained on the Aub* complex together with piRNA precursor transcripts, which are not efficiently processed further. As the interaction between Squ and Spn-E has been predicted by AlphaFold program and verified by the mutational analysis (Kawaguchi et al., 2025), and importantly, the predicted Squ-binding interface on Spn-E is located on the surface opposite to the Aub*-binding interface (Fig. S7C), suggesting that Squ does not directly compete with the Spn-E–Aub* interaction.

**Figure 7:**
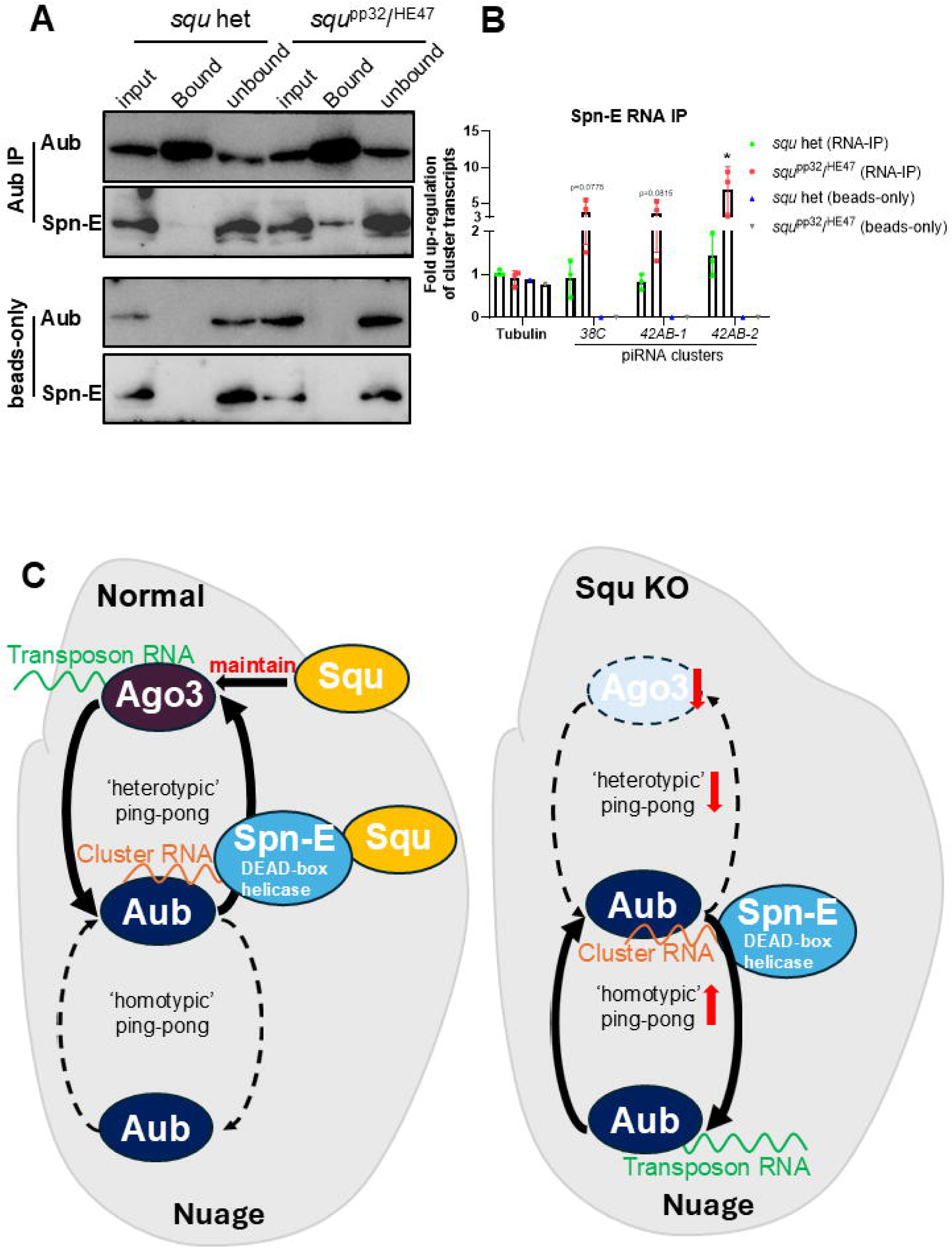
The effect of *squ* loss on Spn-E functions. (A) Immunoprecipitation of Aub in the heterozygous control or *squ*^PP32/HE47^ mutant ovaries. Immunoprecipitation without anti-Aub antibodies is shown as a negative control. (B) RT-qPCR measurement of Spn-E-bound piRNA cluster transcripts from *38C* and *42AB*, and the control *tubulin*. RNAs extracted from Spn-E immunoprecipitates from the heterozygous control or *squ*^PP32/HE47^ mutant ovaries are analyzed. Relative values are obtained by ^ΔΔ^Ct using *rp49* as reference. The signal is further normalized by the Spn-E protein levels in IP fraction (Fig. S7B). Error bars indicate standard deviation (3 biological replicates). Beads-only samples are included as a negative control. Statistics; unpaired T-test (*p <0.05). (C) Model for Squ functions in piRNA biogenesis in nuage. Squ contributes through two mechanisms of Squ: (1) Squ directly binds Spn-E, ensuring efficient heterotypic ping-pong between Aub and Ago3 and promoting processing of interacting transcripts including piRNA precursors. (2) Squ stabilizes Ago3 by unknown mechanisms, supporting proper piRNA production. Upon loss of *squ* function, Spn-E remains bound to the Aub* complex at nuage together with the cleaved transcripts which are normally transferred to Ago3. Reduced Ago3 protein is not efficient for proper heterotypic ping-pong, leading a part of Aub to substitute for Ago3 and engage Aub-Aub homotypic ping-pong, which is not effective for piRNA production and TE repression.

As an RNA helicase protein, Spn-E is expected to possess RNA-binding and ATPase activities (Ott et al., 2014), so we next examined whether Squ directly affects the ATPase activity of Spn-E. Purified wild-type Spn-E protein (Spn-E^WT^) exhibited basal ATP hydrolysis activity *in vitro*, whereas the ATPase-dead mutant, Spn-E^DQVH^, showed no detectable ATPase activity (Fig. S7D). Squ protein alone also did not exhibit ATPase activity (Fig. S7D). Notably, the addition of Squ did not significantly alter the ATP hydrolysis activity of Spn-E^WT^ in these *in vitro* conditions (Fig. S7D). Together, these results demonstrate that, although Squ does not disrupt Aub*–Spn-E binding directly and affect the basal ATPase activity of Spn-E *in vitro*, loss of Squ leads to persistent retention of Spn-E on the Aub* complex together with piRNA precursor transcripts *in vivo*. This aberrant accumulation of the Aub*–Spn-E–RNA intermediate provides a mechanistic explanation for why piRNA precursor processing and downstream piRNA pathway function remain defective in conditions where Ago3 protein levels can be partially restored by Squ in the *spn-E* mutant background.

## Discussion

The piRNA pathway, which represses transposable elements, is well conserved across virtually all animals examined to date, including *Drosophila melanogaster*. In *Drosophila* germline cells, piRNA amplification occurs in the membrane-less perinuclear organelle known as nuage, where multiple piRNA pathway components localize (Aravin et al., 2006; Hirano et al., 2014; Lim and Kai, 2015; Suyama and Kai, 2024). However, the precise mechanisms underlying piRNA biogenesis and the molecular functions of individual nuage components remain elusive. In this study, we systematically characterized the molecular role of Squ, a *Drosophila*-specific nuage protein, and uncovered its dual functions in piRNA pathway: facilitating Spn-E–dependent RNA processing and maintaining Ago3 protein stability (Fig. 7C).

Our results suggest that Squ functions as a co-factor of Spn-E, an RNA helicase and facilitates the efficient transfer of sliced RNA intermediates from Aub to Ago3 during the ping-pong amplification cycle. RNA helicases generally utilize their ATPase activity to unwind RNA structures or release a single-stranded RNA from the duplexes (Jankowsky and Fairman, 2007; Jankowsky, 2011). Considering this general property, Spn-E–dependent RNA remodeling is likely required for efficient heterotypic ping-pong between Aub and Ago3. In support of this concept, the ATPase activity of another nuage-localized RNA helicase, Vas, is stimulated by the nuage factor Tejas (Tej) through its Lotus domain in the presence of single-stranded RNA (Xiol et al., 2014; Jeske et al., 2017). Notably, Tej, like Squ, also binds Spn-E (Lin et al., 2023), suggesting that multiple nuage factors may cooperate to regulate Spn-E activity. In this study, we detected the intrinsic ATPase activity of Spn-E *in vitro*; however, Squ did not significantly enhance Spn-E ATPase activity (Fig. S7D). We speculate that Tej and/or other yet-unidentified Spn-E interacting proteins may serve as primary activators of Spn-E *in vivo*, and these may associate with the Aub*–Spn-E–Squ complex (Fig. S7C). Further experiments are still necessary to elucidate the enzymatic activity of Spn-E and its functional relevance *in vivo*.

Previously, we reported that Squ and Spn-E expressed in S2 cells, a somatic cell line that lacks germline-specific piRNA factors, can directly bind each other. Four interacting residue-pairs between Squ and Spn-E are conserved in *Drosophila* species and mutation of four conserved interacting residues in Squ (Squ^4A^) abolishes this interaction in S2 cells (Kawaguchi et al., 2025). In this study, we further confirmed that Squ^4A^ also fails to properly interact with Spn-E *in vivo* using transgenic expression in the ovary, and that reciprocal mutation of the corresponding residues in Spn-E (Spn-E^4A^) similarly abolishes its interaction with Squ in S2 culture cells (Fig. 5B; Fig. S5B). Importantly, Squ^4A^ failed to rescue the piRNA-deficient phenotypes of the *squ* mutant ovaries (Fig. 5 and S5). Nevertheless, a substantial fraction of Squ^4A^ remained detectable in the perinuclear nuage region (Fig. 5C). A similar discrepancy between physical interaction and nuage localization has been reported for Tej: although Tej-^Δ^SRS loses its ability to interact with Spn-E in S2 culture cells, it does not disrupt the nuage localization of Spn-E or Vas *in vivo*, whereas a bit larger, extended SRS deletion is required to abolish their interaction (Lin et al., 2023). These observations suggest that the highly condensed molecular environment of nuage and/or additional protein interactions can maintain nuage localization even when direct interactions with Spn-E are weakened or disrupted.

Consistent with the reduced Ago3 protein level in *squ* mutant, heterotypic ping-pong between Aub and Ago3 was severely compromised (Fig. 4), whereas Aub protein level was not affected (Fig. 1C). Because Aub–Aub homotypic ping-pong activity was clearly detected in *squ* mutant ovaries, we propose that, when Ago3 is non-functional, sliced RNAs are transferred from one Aub molecule to another to sustain this alternative cycle (Fig. 7C). This homotypic pathway, although active, is less efficient in piRNA production and transposon repression than the heterotypic pathway. Aub–Aub homotypic ping-pong has also been reported in *krimp* and *qin*, also known as *kumo*, mutant ovaries; Krimp promotes Aub–Ago3 coupling through its Tudor-domain–mediated dimerization, and Qin/Kumo is likewise required for Aub–Ago3 association (Zhang et al., 2011; Anand and Kai, 2012; Sato et al., 2015). These observations suggest that efficient heterotypic ping-pong is supported by the coordinated actions of multiple regulatory factors, whereas the Aub–Aub cycle can become prominent when this regulatory network is perturbed. In this study, we show that Squ as one such regulatory factor that promotes heterotypic ping-pong through its functional coupling with Spn-E (Fig. 7C).

Our results demonstrate that Squ is essential for maintaining Ago3 protein stability. Loss of Squ led to a marked reduction of Ago3 protein level and Ago3-bound piRNAs, while *ago3* transcript level was not affected (Fig. 1E-F, 3A). These results suggest that Squ regulates Ago3 via a post-transcriptional mechanism. It has been reported that unloaded Piwi protein is destabilized and degraded (Sienski et al., 2012; Kina et al., 2025), and we therefore propose that the piRNA-free, ‘‘empty’’ form of Ago3 may likewise be subjected to proteasome-dependent turnover. However, ectopic expression of Squ in *spn-E* mutant ovaries restored Ago3 protein abundance despite the severe reduction in piRNA production caused by loss of *spn-E* function, implying that Squ may possess an additional, direct role on stabilizing Ago3 protein (Fig. 6). Notably, restoration of Ago3 protein level in *spn-E* mutant did not rescue piRNA precursor processing defects and transposon de-repression, indicating that *spn-E* is genetically epistatic to *squ* and *ago3* function. These results uncover a functional hierarchy among nuage components, where Squ contributes to transposon repression by supporting both Spn-E–mediated precursor processing and Ago3 stability via an unresolved post-transcriptional mechanism.

Taking together, our results refine the current model of piRNA biogenesis in *Drosophila* germline cells. Squ functions on stabilizing Ago3 protein for heterotypic ping-pong between Aub and Ago3. Squ also acts as a crucial co-factor of Spn-E in the nuage, promoting piRNA precursor turnover or facilitating transfer of sliced transcripts between ping-pong partners. Further studies are needed to elucidate the molecular mechanism for Squ to perform these functions. Understanding these mechanisms will provide deeper insights into the piRNA production mechanism cooperated by many regulatory proteins.

## Materials and Methods

### Fly stocks

All fly stocks were maintained at 25°C on standard food (5% (w/v) dry yeast, 5% (w/v) corn flower, 2% (w/v) rice bran, 10% (w/v) glucose, 0.7% (w/v) agar, 0.2% (v/v) propionic acid, and 0.05% (w/v) p-hydroxy butyl benzoic acid). The mutant alleles used in this study were *squ*^PP32/HE47^ (Pane et al., 2007), *ago3*^g1/g2^ (VDRC #313531; VDRC #313532; Senti et al., 2015), *spn-E*^7-4^ (this study), *tej*^48-5^/Df(2R) Exel7131 (Patil and Kai, 2010; BL #7876), *vas*^4-2^ (this study) and *krimp*^f06583^/Df(2R) Exel6063 (BL #18990, Lim and Kai, 2007; BL #7545). The heterozygotes of individual mutants were used as the control conditions. Knock-In fly line *aub*^EGFP.KI^ (DGRC #118621, Kina et al., 2019) and transgenic fly line UASp-3xFlag.AGO3.WT (DGRC #118699) were obtained from the *Drosophila* Genetic Resource Center at the Kyoto Institute of Technology, Japan. *tej*^EGFP.KI^ was generated in our laboratory (Lin et al., 2023). NGT40-Gal4; *nos*-Gal4-VP16 (Grieder et al., 2000) was used as germline driver line. The genotypes of all fly strains are listed in Table S1.

### Generation of knock-out flies

Loss of function alleles, *spn-E*^7-4^ and *vas*^4-2^, were generated using the CRISPR/Cas9 system. Guide RNAs (gRNAs) targeting the *spn-E* and *vas* loci were designed and cloned into pDCC6 vector, respectively (Gokcezade et al., 2014). The sequences of the gRNAs were: Spn-E gRNA: 5’ -GTGCCCAACCGCGAGTTGAT-3’ and Vasa gRNA: 5’-GGGGATACAATCACTACCTG-3’. Obtained gRNA expression plasmids (120 ng/μl) were injected into *y w* embryos. PCR-amplified genomic fragments from candidate lines were subjected to Sanger sequencing, which revealed that *spn-E*^7-4^ contains an 8-bp deletion, GCGAGTTG, at 15837616-15837623 (release=r6.53), together with a 4-bp insertion, AACA, causing a frameshift and premature stop codon. Similarly, *vas*^4-2^ carries a 4-bp deletion, GTAG, at 15072945-15072948 (release=r6.53), causing a frameshift leading to premature termination. Loss of the corresponding protein products was confirmed by western blotting and immunostaining using anti-Spn-E and anti-Vas antibodies, respectively.

### Generation of transgenic flies

The transgenic fly lines carrying native squ promoter-GFP-3xFlag-Squ^WT^ (pNative-Squ^WT^) or -Squ^4A^ (pNative-Squ^4A^) were generated using PhiC31-mediated integration system. The DNA fragments encoding Squ^WT^ or Squ^4A^ were amplified from pENTR-Squ^WT^ and Squ^4A^ plasmids, respectively (Kawaguchi et al., 2025). The fragments of promoter region, 5’UTR and 3’UTR were amplified by PCR from genomic DNA. GFP-3xFlag fragment was amplified from the plasmid containing GFP-3xFlag-tag sequence. All the fragments are combined and cloned into pUASt-attB vector (Brand and Perrimon, 1993) by the reaction of In-Fusion® Snap Assembly Master Mix (Takara). The generated constructs were introduced into attP2 site (BDSC#25710).

### Antibodies

Polyclonal rabbit anti-mKate2 antibody was generated in this study. His-tagged full-length mKate2 was expressed in *Escherichia coli* BL21(DE3) from the pDEST17 (Thermo Fisher Scientific) plasmid carrying the mkate2 coding sequence. His-tagged mkate2 protein was solubilized with 6 M urea in PBS and purified using Nickel Sepharose beads (GE Healthcare) according to the manufacturer’s protocol. The antibodies used in this study were rat anti-Squ (1:1000, Kawaguchi et al., 2025), guinea pig anti-Aub (1:1000, Lim et al., 2022), mouse monoclonal anti-Ago3 (1:500, National Institute of Genetics, Japan), guinea pig anti-Vas (1:2000, Patil and Kai, 2010), rat anti-Spn-E (1:500, Patil and Kai, 2010), rat anti-Tej (1:500, Lin et al., 2023), guinea pig anti-Krimp (1:1000, Lim and Kai, 2007), guinea pig anti-Ago2 (1:100, Lin et al., 2023), rabbit anti-mkate2 (1:1000, this study), rabbit anti-HeT-A Gag polyclonal antibody (1:1000, Lin et al., 2023), mouse monoclonal anti-Tubulin (1:1000, DM1A, sc-32293, Santa Cruz Biotechnology). The secondary antibodies used in this study were HRP-conjugated goat anti-guinea pig (1:1000, Dako, Cat. #P0141), HRP-conjugated goat anti-rat (1:1000, Dako, Cat. # P0450), HRP-conjugated goat anti-mouse (1:3000, BioRad, Cat. # 1706516) and HRP-conjugated goat anti-rabbit (1:3000, BioRad, Cat. #1706515). ANTI-FLAG^®^ M2-Peroxidase (1:1000, Sigma, Cat. #A8592) and HRP-conjugated anti-Myc-tag antibody (1:1000, MBL, Cat. # M192-7) were also used.

### Western blotting

Ovaries were homogenized in the ice-cold PBS and denatured in the presence of 1×SDS sample buffer at 95°C for 5 min. The protein samples were subjected to SDS-PAGE and transferred to ClearTrans SP PVDF membrane (Wako). Primary and secondary antibodies described above were diluted in the Signal Enhancer reagent HIKARI (Nacalai Tesque). Chemiluminescence was induced by the Chemi-Lumi One reagent kit (Nacalai Tesque) and detected by ChemiDoc Touch (Bio-Rad). The immunoblotting signals were quantified using ImageJ (Schneider et al., 2012) or Image Lab software (Bio-Rad).

### RNA immunoprecipitation

To analyze RNAs interacting with proteins, we followed the methods described previously (Lin et al., 2023). To extract Aub- and Ago3-bound piRNAs, and Spn-E–bound RNAs, 300 ovaries (Aub), 500 ovaries (Ago3) and 400 ovaries (mKate2-Spn-E) were dissected from 1-day-adult females in cold PBS buffer. Ovaries were homogenized in lysis buffer containing 20 mM Tris-HCl (pH 7.4), 200 mM NaCl, 2 mM DTT, 10% (v/v) glycerol, 2 mM MgCl_2_, 1% (v/v) Triton X-100, 1× cOmplete protease inhibitor cocktail (Roche), and 1% (v/v) RNaseOUT recombinant ribonuclease inhibitor (Invitrogen). The lysates were cleared by centrifugation at 20,000 ×*g* for 10 min at 4°C three times to remove cell debris. Dynabeads Protein G/A (Invitrogen) pre-equilibrated with lysis buffer were added to the cleared lysates and rotated at 4°C for 1 h for pre-clearing. After clearance, guinea pig anti-Aub (1:10, Lim et al., 2022) or mouse monoclonal anti-Ago3 (1:500, NIG) or rabbit anti-mKate2 (1:500, this study) was added to the lysates and incubated for 3 h with gentle rotation at 4°C. Dynabeads Protein G/A were then added to the mixture and incubated at 4°C for 1 h with rotation. After incubation, magnet beads were collected and washed 5 times by wash buffer containing 20 mM Tris-HCl (pH7.4), 400 mM NaCl, 2 mM DTT, 10% (v/v) glycerol, 2 mM MgCl_2_, 1% (v/v) Triton X-100, 1× cOmplete protease inhibitor cocktail (Roche), and 1% (v/v) RNaseOUT recombinant ribonuclease inhibitor (Invitrogen). After washing, beads were resuspended in 100 μl wash buffer. Ten μl was used for western blotting to check the efficiency of protein immunoprecipitation and the remaining 90 μl solution was used for RNA extraction using TRIzol LS (Invitrogen) following standard manufacturer’s protocol. Extracted RNA were stored at -80°C until analysis. For mKate2-Spn-E RNA IP, negative control was performed by adding equal amount of Dynabeads Protein G/A with IP groups but without antibody.

### Small RNA sequencing and analysis

To compare Aub- and Ago3-bound piRNAs in two conditions, the amount of Aub or Ago3 proteins in IP samples were equalized using the signals in the immunoblotting analysis. Libraries from Aub- and Ago3-bound RNAs were prepared and sequenced on an Illumina HiSeq 2500 platform (Genome Information Research Center, Research Institute for Microbial Diseases, Osaka University). After adaptor trimming (5’-AGATCGGAAGAGCACACGTCT-3’), reads mapping to miRNA, rRNA, snRNA, snoRNA and tRNA sequences were removed. Remaining reads were size-selected with 23-29 nt and mapped to major piRNA clusters or the consensus sequences of transposable elements using Bowtie allowing up to 3 mismatches (Langmead et al., 2009). piRNA cluster definition was according to previous paper (Brennecke et al., 2007) and TE sequences were from Flybase (Release 6.32). Read counts were normalized with the counts for 21nt-spike RNA (ACATGGCATGGATGAACTATA) which were added equally to individual RNA samples prepared for library construction (Lutzmayer et al., 2017). piRNA cluster-mapping reads were visualized by pyGenomeTracks (Ramírez et al., 2018). Nucleotide preference was visualized by sequence logo using Ggplot2 R ggseqlogo (Wagih, 2017).

### Crosslinking immunoprecipitation (CL-IP)

200 ovaries were dissected from 1-day-adult *y w* females in ice-cold PBS and fixed in PBS containing 0.1% (w/v) paraformaldehyde for 20 min on ice. Crosslinking was quenched in 125 mM glycine for 20 min at room temperature. Ovaries were homogenized in CL-IP lysis buffer containing 50 mM Tris-HCl (pH 8.5), 150 mM KCl, 5 mM EDTA, 1% (v/v) Triton X-100, 0.1% (w/v) SDS, 0.5 mM DTT, and 1× cOmplete protease inhibitor cocktail. The lysate was incubated at 4°C for 20 min with rotation and then sonicated with Bioruptor (Lin et al., 2023). After centrifugation at 20,000 ×*g* for 10 min at 4°C, the supernatant was collected and diluted with an equal volume of CL-IP wash buffer containing 25 mM Tris-HCl (pH 7.5), 150 mM KCl, 5 mM EDTA, 0.5% (v/v) Triton X-100, 0.5 mM DTT, and 1× cOmplete protease inhibitor cocktail. Pre-washed Dynabeads Protein G/A mixture (1:1) were added for pre-clearance at 4°C for 1 h with rotation. Anti-Squ antibody was added to pre-cleared supernatant with 1:500 dilution and incubated at 4°C overnight. The pre-equilibrated Dynabeads Protein G/A mixture (1:1) were added for binding and incubated at 4°C for 3 h with rotation. After washing three times with CL-IP wash buffer, beads were collected and resuspended in 50 μl of CL-IP wash buffer containing 1× SDS sample buffer. The beads were then boiled at 95°C for 5 min and subjected to SDS-PAGE for western blotting analysis. Negative control was performed as the description above.

### Immunostaining of ovaries

Ovaries were dissected, fixed with 5% (w/v) paraformaldehyde containing PBS for 20 min and permeabilized in PBX (PBS with 0.2% [v/v] Triton X-100). The primary antibodies used in this study were described above. Secondary antibodies were Alexa Fluor 488-, 555-, and 633-conjugated goat antibodies at a 1:200 dilution (Thermo Fisher Scientific, United States) and CF®633 goat antibodies at a 1:1000 dilution (Biotium, Fremont, CA, United States). Ovaries expressing endogenous fluorescent-tagged proteins were fixed with 0.1% (w/v) paraformaldehyde containing PBS for 20 min and washed with PBX for 15 min, twice. DAPI was diluted for 1:1000 to stain DNA. Ovaries were equilibrated in Fluoro-KEEPER Antifade Reagent (Nacalai Tesque) and stored at 4℃ before mounting. Images were taken by ZEISS LSM 900 with Airy Scan 2 using 63x oil NA 1.4 objectives and processed by ZEISS ZEN 3.0 and ImageJ (Schneider et al., 2012).

### RNA in situ hybridization chain reaction (HCR)

Probes (Molecular Instruments, *42AB*-LOT: PRJ124; *38C*-LOT: PRJ125) targeting piRNA cluster transcripts from specific regions of cluster *42AB* (Chr2R: 6322410-6323756) and *38C* (Chr2L: 20104896-20213637) were generated and used in this experiment. Commercial reagents were used for this reaction (Molecular Instruments, Inc.) according to the manufacturer’s instructions and previously described protocols (Lin et al., 2023).

### Quantification of in situ-HCR signal for piRNA precursors

Fluorescence intensities for piRNA clusters *38C* and *42AB* were quantified using ImageJ (Fiji) after background subtraction. WGA was used to stain the nuclear envelope, and a region extending ±5% of the nuclear diameter inside and outside the envelope was defined as the perinuclear region, as previously described (Lin et al., 2023).

### RT-qPCR

Flies were fattened up with moist yeast paste for 2 days and then dissected for RNA extraction using TRIzol LS (Invitrogen) following manufacturer’s protocol with DNase I (Invitrogen) treatment. Reverse-transcription was performed by Super Script III system (Invitrogen) using primers of a mixture for d(T)20 and randomized hexadeoxyribonucleotide. KAPA SYBR Fast qPCR Master Mix (KAPA biosystems) was used for qPCR reactions and primer sequences used in this study were listed in Table S2. The relative expression levels were calculated by ^ΔΔ^Ct method using *rp49* as reference. All the experiments were performed by 3 biological replicates.

### mRNA-seq

Total RNAs were extracted from *squ* heterozygous and mutant ovaries using TRIzol LS (Invitrogen) following the manufacturer’s instruction. Poly(A)+ RNA was enriched using oligo(dT) beads with the NEBNext Poly(A) mRNA Magnetic Isolation Module (New England Biolabs, USA). The enriched RNA was fragmented, reverse-transcribed, adapter-ligated, and PCR-amplified to prepare cDNA libraries using the NEBNext Ultra II Directional RNA Library Prep Kit (New England Biolabs, USA). Library preparation and sequencing were outsourced to Rhelixa (Japan), yielding approximately 35 million reads per sample. Low quality sequences and adapters were removed by fastp (Chen et al., 2018). Paired end reads were mapped to *Drosophila melanogaster* reference genome r6.32 and the canonical sequences of TEs by STAR version 2.7 with default setting. (Dobin et al., 2013). Reads mapped to exons of individual genes and the canonical TEs were counted using featureCounts (Liao et al., 2014). Differential expression analysis was performed by DESeq2 (Love et al., 2014) with significant up-regulation (Log_2_FC>1, p-adj<0.05) and down-regulation (Log_2_FC<−1, p-adj<0.05). Two biological replicates were performed for the mRNA-seq analysis.

### Co-immunoprecipitation in S2 cells

The *Drosophila* S2 cell line (S2-DRSC) was cultured at 28°C in Schneider’s medium supplemented with 10% (v/v) fetal bovine serum and antibiotics (penicillin and streptomycin). Protein-coding sequences of *squ* and *spn-E* were amplified and cloned into the pENTR vector (Thermo Fisher Scientific) and subsequently transferred into pAFW or pAMW destination vectors for plasmid construction. S2 cells (0.2–2 × 10^6^ cells/ml) were pre-cultured in 12-well plates overnight before transfection with Hilymax (Dojindo Molecular Technologies, Japan). After 36–48 h of transfection, S2 cells were collected for co-immunoprecipitation (co-IP) experiments as previously described (Kawaguchi et al., 2025).

### *Drosophila* fertility test

Female fertility was assessed by pairing three *y w* males with five females of each genotype in an enclosed environment on apple juice plates supplemented with a small amount of moist yeast paste. The number of eggs laid was recorded after 24 h of mating, and the hatching rate was calculated 24 h after egg laying. Egg counts and hatching rates were recorded for at least 3 consecutive days.

### Purification of recombinant proteins in Sf9 cells

*Spodoptera frugiperda* Sf9 cells were cultured using ExpiSf^TM^ CD Medium (Thermo Fisher Scientific/Gibco) in 27℃ on an orbital shaker platform at 120 rpm. The fragment of DNA product for 6His-2StrepII-GFP-Spn-E^WT^, -Spn-E^DQVH^ or -Squ^WT^ was amplified and inserted into pFastBac1 vector (Thermo Fisher Scientific). Recombinant baculoviruses expressing 6His-2StrepII-GFP–Spn-E^WT^, –Spn-E^DQVH^, or –Squ^WT^ were generated using the NucleoBond® Xtra BAC kit (Clontech/Takara) and used to infect Sf9 cells according to the manufacturer’s protocol (Thermo Fisher Scientific). After several days, infected Sf9 cells were resuspended in 10 ml lysis buffer: 100 mM Tris-HCl (pH 8.0), 150 mM NaCl, 1% TritonX-100, 1 mM DTT, 1x cOmplete protease inhibitor cocktail (PIC, Roche) and 1 mM PMSF, sonicated for total 10 min with 1 min on/1 min off on the ice and centrifuged for 1 h at 15,000 ×g to collect the supernatant. 1 ml Strep-Tactin® Sepharose® resin beads (IBA Lifesciences) were added to the Poly-Prep Chromatography Column (Bio-rad) and followed with 10 ml wash buffer: 100 mM Tris-HCl (pH 8.0), 500 mM NaCl, 1 mM DTT and 1 mM PMSF. All supernatant was then loaded into columns and washed with 10 ml wash buffer, protein was eluted by 10 ml elution buffer: 100 mM Tris-HCl (pH 8.0), 150 mM NaCl, 1 mM DTT, 5 mM desthiobiotin and 1x PIC. Recombinant proteins were concentrated by vivaspin turbo 15 (Nippon Genetics Co., Ltd) and stored in -80℃ after rapid freezing by liquid nitrogen. SDS-PAGE and CBB staining were performed to check the purification and quantity for protein.

### ATPase assay

ATP hydrolysis reactions were performed at 25°C for 1, 2, 3, or 5 h with 25 μM recombinant protein (6His-2StrepII-GFP-Spn-E^WT^, Spn-E^DQVH^, or Squ^WT^) in ATPase assay buffer (10 mM Tris-HCl, pH 8.0, 100 mM NaCl, 2 mM MgCl_2_, 1 mM DTT, and 200 μM ATP (Sigma)). To examine the effect of Squ on Spn-E ATPase activity, Squ protein was added at 25 µM to the Spn-E ATPase reaction mixtures. Inorganic phosphate was quantified using BIOMOL® Green Reagent (Enzo Life Sciences) in 96-well plates according to the manufacturer’s protocol. Absorbance at 620 nm was measured with an SH 9000 microplate reader (Corona Electric), background values were subtracted, and phosphate concentrations were normalized to calculate ATP hydrolysis (%) using a phosphate standard curve.

## Acknowledgments

We are grateful to Wakana Isshiki (The University of Osaka) for generation of Squ antibody, to Haitian Qin (The University of Osaka) for assisting of generation of mKate2 antibody, to Dr. Hilo Yen (The University of Osaka) for generation of *spn-E* and *vas* mutant alleles, to Dr. Ritsuko Suyama (The University of Osaka) for HCR-FISH experiments and data analysis. We are also grateful to Dr Trudi Schüpbach (Princeton University) for generous gifts of *squ* mutant flies. We acknowledge Bloomington *Drosophila* Stock Centre and Kyoto Stock Center for the fly stocks. We also thank for the FBS Core Facility in The University of Osaka for providing access to the LSM 900 and ChemiDoc Touch. We thank for the Center for Medical Research and Education, Graduate School of Medicine, The University of Osaka for microplate reader SH-9000Lab (Corona Electric, Japan). We appreciate the insightful discussion and suggestions from all the members of TK’s laboratory.

## Author contribution

Conceptualization: X. Xu, S. Kawaguchi, and T. Kai.

Data Curation: X. Xu and S. Kawaguchi.

Formal Analysis: X. Xu and S. Kawaguchi.

Funding Acquisition: S. Kawaguchi and T. Kai.

Investigation: X. Xu.

Methodology: X. Xu, S. Kawaguchi, T. Iki, and T. Kai.

Project Administration: T. Kai.

Resources: S. Kawaguchi, T. Iki and T. Kai.

Software: X. Xu, T. Iki and S. Kawaguchi.

Supervision: S. Kawaguchi, T. Iki and T. Kai.

Validation: X. Xu and S. Kawaguchi.

Writing, Original Draft: X. Xu and S. Kawaguchi.

Writing, Review and Editing: X. Xu, S. Kawaguchi, T. Iki and T. Kai.

## Data Availability

NGS data set have been deposited to the DNA Data Bank of Japan (DDBJ) and under BioProject ID PRJDB37636.

## Funding

This work was supported by Grant-in-Aid for Transformative Research Areas (A) [21H05275 to T.K.], Grants-in-Aid for Scientific Research [T25K022050 for T.K.], and The University of Osaka D3 Center “Transdisciplinary Research Project” [Na22990007 to S.K.]. X.X. is supported by JST SPRING [J219713005].

## Supplemental information

**Figure S1: Subcellular localization and expression of Squ in *tej* and *vas* mutants**

(A–B) Protein levels and immunostaining of Squ in control ovaries and in *tej* (A) or *vas*

(B) mutant egg chambers. Scale bar is 5 μm.

**Figure S2:**

(A) The heatmap of fold changes of TEs’ expression for all 119 canonical transposons in control and *squ* mutant ovaries.

**Figure S3: Second replicate for Aub- and Ago3- bound piRNAs in the control and *squ* mutant**

(A) Aub- (upper panels) and Ago3-bound (lower panels) small RNAs from control and *squ* mutan ovaries mapped to the major piRNA clusters *38C-3* and *42AB* (replicate 2). Plus-strand (blue) and minus-strand (red) piRNAs are shown as upward and downward peaks, respectively.

(B) Antisense bias of Aub-bound piRNAs mapping to transposable elements (TEs) in control (*squ* het) and *squ* mutant ovaries (replicate 2). Each dot represents one TE; antisense piRNA ratios are calculated by dividing antisense-mapping piRNA reads by total mapping reads for each TE. Horizontal lines indicate the mean values. ****p < 0.0001 (unpaired t-test).

(C) Effect of loss of *squ* function on antisense ratio of Aub-bound piRNAs mapping to transposon consensus sequences. Dots indicate 24 up-regulated TEs (Fig. 2A) in *squ* mutant relative to control ovaries (replicate 2). For each TE, the antisense bias in *squ* mutant is divided by that in *squ* heterozygous.

(D) RT–qPCR analysis of transcripts derived from major germline piRNA clusters *38C* and *42AB*, with tubulin as a control. All values are normalized to *rp49* and are shown relative to expression levels in the presence of *squ* heterozygous. Error bars indicate standard deviation (3 biological replicates). Statistical significance is assessed by Student’s unpaired t-test.

**Figure S4: Second replicate for Aub-Aub homotypic signature in *squ* and *ago3* mutants**

(A) Nucleotide bias of Aub-bound and Ago3-bound piRNAs mapped to piRNA cluster loci in control and *squ* mutant ovaries (replicate 2). Sequence logos indicate 1^st^ to 20^th^ of piRNAs. 10A signature of *squ* mutant condition reflects enhanced Aub–Aub homotypic ping-pong. NA (Not analyzed); Ago3-bound piRNAs are not detectable in *squ* mutant.

(B) Nucleotide bias of Aub-bound piRNAs in control and *ago3* mutant ovaries (replicate 2).

(C) Aub-bound piRNAs from control and *ago3* mutant ovaries mapped to the major germline piRNA clusters *38C-3* and *42AB*. Blue; genomic plus-strand reads, red; minus-strand reads.

(D) Antisense bias of Aub-bound piRNAs mapping to transposable elements (TEs) in control and *ago3* mutant ovaries (replicate 2). Each dot represents one TE; antisense piRNA ratios are calculated by dividing antisense-mapping piRNA reads by total mapping reads for each TE. Horizontal lines indicate the mean values. ****p < 0.0001 (unpaired t-test).

(E) Ratio of antisense bias of Aub-bound piRNAs for the 24 up-regulated TEs (detected in *squ* mutant) in *ago3* mutant versus control ovaries (replicate 2). Dots indicate 24 up-regulated TEs (Fig. 2A) in *ago3* mutant relative to control ovaries (replicate 2). For each TE, the antisense bias in *ago3* mutant is divided by that in *ago3* heterozygous.

**Figure S5: Interaction of Squ and Spn-E is crucial for *Drosophila* female fertility**

(A) Localization of Squ, Ago3 and Krimp in the *squ* heterozygous control germline cells. (upper panels: stage 4–5 egg chambers; lower panels: magnified nuclei indicated by squares in the upper panels). Scale bar is 5 μm.

(B) Immunoprecipitation of Myc-tagged Spn-E^WT^ and Spn-E^4A^ using anti-Myc antibodies with the co-expression of Flag-Squ in S2 culture cells. Flag-Squ can be detected in Spn-E^WT^ but not in Spn-E^4A^ immunoprecipitates.

(C) Immunoblotting of proteins extracted from the heterozygous control, *squ*^PP32/HE47^ and *squ*^PP32/HE47^expressing Squ^WT^ or Squ^4A^ ovaries.

(D) RT–qPCR analysis of transcripts derived from major germline piRNA clusters *38C* and *42AB*, with tubulin as a control. All values are normalized to *rp49* and are shown relative to expression levels in *squ*^PP32/HE47^; pNative-Squ^WT^ ovaries. Error bars indicate standard deviation (3 biological replicates). Statistical significance is assessed by Student’s unpaired t-test. *p <0.05.

(E) Analysis of egg laying and hatching rate. Infertility of *squ* mutant is rescued by the pNative-promoter-Squ^WT^ but not Squ^4A^ mutant. Daily egg production by five females is recorded (3 biological replicates). T-test is performed for the statistics analysis. (****p <0.0001).

**Figure S6:**

(A) Immunoprecipitation of GFP–3×Flag–Squ and the variants expressed under the native promoter (pNative-Squ^WT^ and pNative-Squ^4A^) using anti-3FLAG antibodies in the ovaries expressing pNative-Squ^WT^ and - Squ^4A^ under the loss of endogenous *spn-E* condition.

(B) Immunoblotting of proteins extracted from the heterozygous control, *squ*^PP32/HE47^, *squ* heterozygous and *squ*^PP32/HE47^ expressing NGT40; nosGal4 driven Flag-Ago3 protein. Two bands are detected using anti-Ago3 antibodies (upper bands are the NGT40; nosGal4 driven Flag-Ago3 and lower bands are the endogenous Ago3).

**Figure S7: Structural modeling and in vitro characterization of Spn-E and Squ**

(A) Quantification of intensity ratio for Aub–Spn-E interaction in the presence and absence of *squ*. Relative values are obtained by ^ΔΔ^Ct using *rp49* as reference. The signal is further normalized by the Aub protein levels in IP fractions. Error bars indicate standard deviation (3 biological replicates). The beads only controls are also shown as a negative control. Statistics; unpaired T-test.

(B) Immunoblotting of the input and Spn-E IP fraction samples in the presence and absence of *squ*. 2, 4, 8 and 16 μl of the IP samples are loaded to perform the comparison of IP efficiency.

(C) Predicted structure of the SIWI*/Aub* complex with Spn-E and Squ.

(i) Cryo-EM structure of the *Bombyx mori* SIWI/Aub–piRNA–target RNA–GTSF1–Maelstrom–Spn-E complex (PDB: 9NHS). SIWI is shown in gray, GTSF1 in orange, Maelstrom in purple, and Spn-E in dark green. The target RNA is depicted as a cyan ribbon.

(ii) Predicted *Drosophila melanogaster* Spn-E–Squ complex generated using AlphaFold3. Spn-E is shown in green and Squ in yellow. A single-stranded RNA is displayed as a blue ribbon, and ATP is shown in stick representation.

(iii) Structural superposition of the two models based on alignment of their Spn-E subunits.

(D) ATPase activity of purified Spn-E^WT^, Spn-E^DQVH^, and Squ. ATP hydrolysis is measured up to 5 h, and released inorganic phosphate is quantified. Spn-E^WT^ shows robust ATPase activity, whereas Spn-E^DQVH^ and Squ alone exhibit minimal ATP hydrolysis. Error bars indicate standard deviation (3 biological replicates). Statistics; unpaired T-test (***p <0.001).

## Notes

### Competing Interest Statement

The authors have declared no competing interest.

### Summary of Updates

1. The mistake of the labeling of the legends in Figrue 3D for the 'yellow' has been revised to 'white'. 2. The mistake of the labeling of the legends in Figrue S5A for the 'squ heterozygous control and mutant germline cells' has been revised to 'squ heterozygous control germline cells'.

