## Supplementary material for "Squash ensures Spindle-E–dependent heterotypic ping-pong amplification of piRNAs in the *Drosophila* ovary": Sup. Table1

**Supplementary Table 1. *Drosophila* genotypes used in this study.**

| **Genotypes** | **Source** |
| --- | --- |
| *w^[-]^; squ^pp32^/CyO; TM3, Sb/ TM6, Tb* | Gift from (Pane et al., 2007) |
| *w^[-]^; squ^HE47^/CyO; TM3, Sb/ TM6, Tb* | Gift from (Pane et al., 2007) |
| *w^[-]^;Pin/CyO; spn-E^7-4^/TM3, Sb* | This study |
| *w^[-]^; tej^48-5^*/*CyO* | Patil and Kai, 2010 |
| *w^[-]^; Df(2R)Exel7131/CyO* (Df stock for *tej* mutant) | BL#7876 |
| *w^[-]^; vas*^4-2^*/CyO* | This study |
| *w^[-]^; PBac{WH}krimp^f06583^/CyO* | BL #18990  (Lim and Kai, 2007) |
| *w^[-]^; Df(2R)Exel6063/CyO* (Df stock for *krimp* mutant) | BL#7545 |
| *w^[-]^; ago3^g1^/TM3,Sb* | VDRC#313531 |
| *w^[-]^; ago3^g2^/TM3,Sb* | VDRC#313532 |
| *w^[-]^; squ^HE47^/CyO;* native-promoter-GFP-3xFlag-Squ^WT^/TM3, Sb | This study |
| *w^[-]^; squ^HE47^/CyO;* native-promoter-GFP-3xFlag-Squ^4A^/TM3, Sb | This study |
| *w^[-]^; Pin/CyO; spn-E^7-4^,* native-promoter-GFP-3xFlag-Squ^WT^*/TM3, Sb* | This study |
| *w^[-]^; Pin/CyO; spn-E^7-4^,* native-promoter-GFP-3xFlag-Squ^4A^*/TM3, Sb* | This study |
| *w^[-]^; squ^pp32^, NGT40-*Gal4/ *CyO; nos-*Gal4 VP16 | This study |
| *w^[-]^; squ^HE47^*/*CyO;* UASp-3xFlag.AGO3.WT / *TM3, Sb* | This study |
| *w^[-]^; squ^pp32^/CyO;* Spn-E-mk2 | This study |
