## Supplementary material for "Squash ensures Spindle-E–dependent heterotypic ping-pong amplification of piRNAs in the *Drosophila* ovary": Sup. Table2

**Supplementary Table 2. List of primers used for qRT-PCR in this study.**

| Primer label | Primer sequence | Source |
| --- | --- | --- |
| Rp49_Fw | ATGACCATCCGCCCAGCATAC | (Lin et al., 2023) |
| Rp49_Rv | CTGCATGAGCAGGACCTCCAG | (Lin et al., 2023) |
| Tubulin_Fw | GGTAACCGTCGAAATCAGTGTT | (Lin et al., 2023) |
| Tubulin_Rv | TGGCTTTTCTGCTATACGTGTC | (Lin et al., 2023) |
| Actin5c_Fw | AAGTTGCTGCTCTGGTTGTCG | **Com E9 where from?** |
| Actin5c_Rv | GCCACACGCAGCTCATTGTAG | **Com F2 where from?** |
| 38C_Fw | TCCGTGACGGTTTAGCCCA | (Lin et al., 2023) |
| 38C_Rv | AGGTTTCAAACCTTCCAG | (Lin et al., 2023) |
| 42AB #1_Fw | CGTCCCAGCCTACCTAGTCA | (ElMaghraby et al., 2019) |
| 42AB #1_Rv | ACTTCCCGGTGAAGACTCCT | (ElMaghraby et al., 2019) |
| 42AB #2_Fw | CGCTGTTGAAAGCAAATTGA | (ElMaghraby et al., 2019) |
| 42AB #2_Rv | GAGACCTTCGCTCCAGTGTC | (ElMaghraby et al., 2019) |
| blood_Fw | CCAACAAAGAGGCAAGACCG | (Zhang et al., 2021) |
| blood_Rv | TCGAGCTGCTTACGCATACTGTC | (Zhang et al., 2021) |
| HMS-Beagle_Fw | ctgttgacccttattcgccg | This study |
| HMS-Beagle_Rv | ccgagaatccgagcactttg | This study |
| Burdock_Fw | AGGGAAATATTTGGCCATCC | (Czech et al., 2013) |
| Burdock_Rv | TTTTGGCCCTGTAAACCTTG | (Czech et al., 2013) |
