## Supplementary figures and images for "Squash ensures Spindle-E–dependent heterotypic ping-pong amplification of piRNAs in the *Drosophila* ovary"

### Sup.Figure1

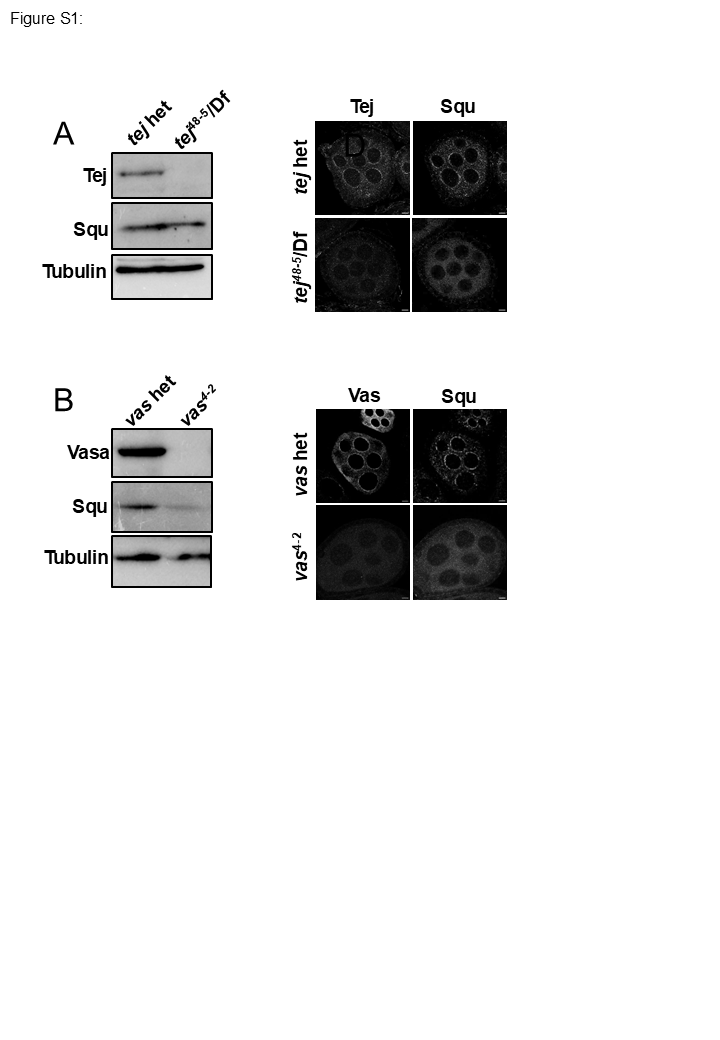

### Sup.Figure2

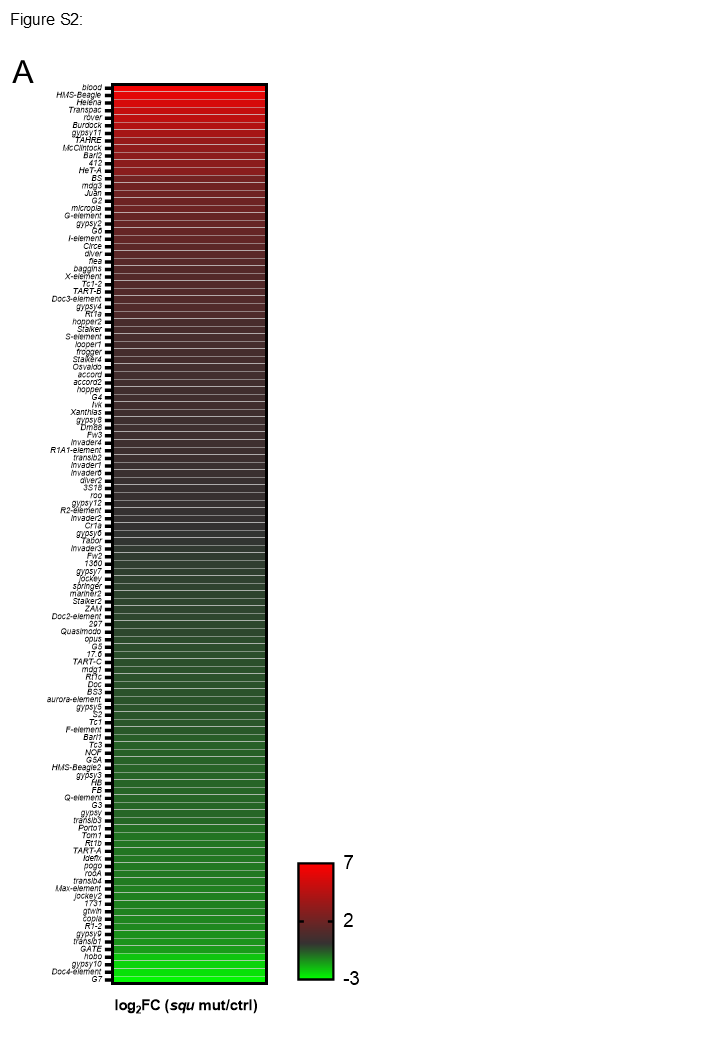

### Sup.Figure3

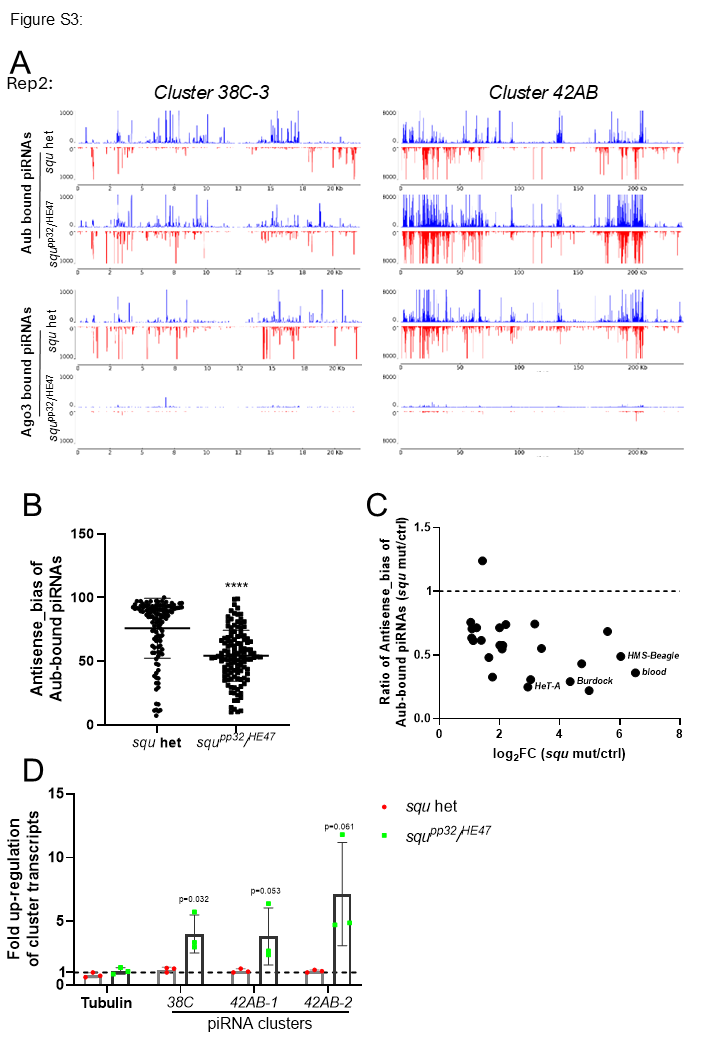

### Sup.Figure4

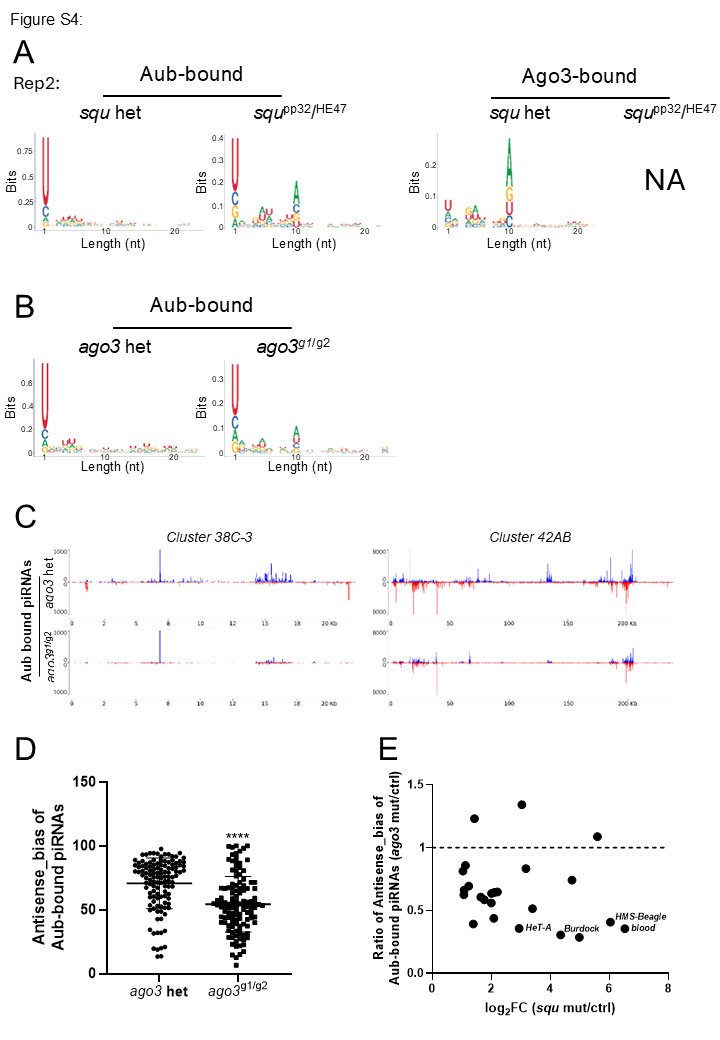

### Sup.Figure5

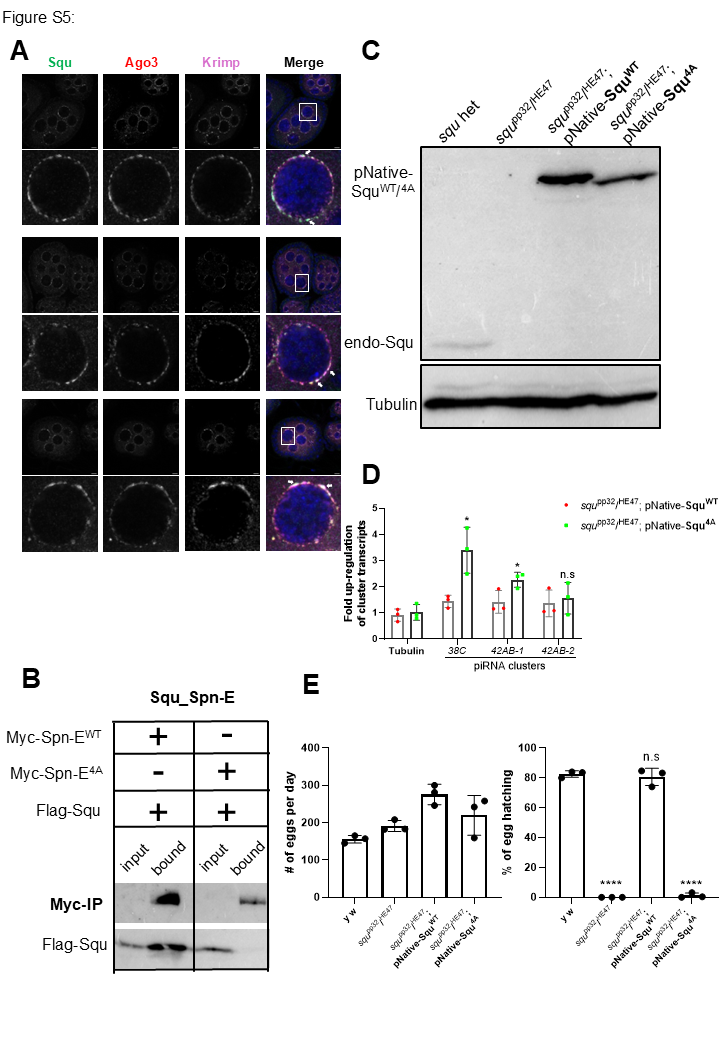

### Sup.Figure6

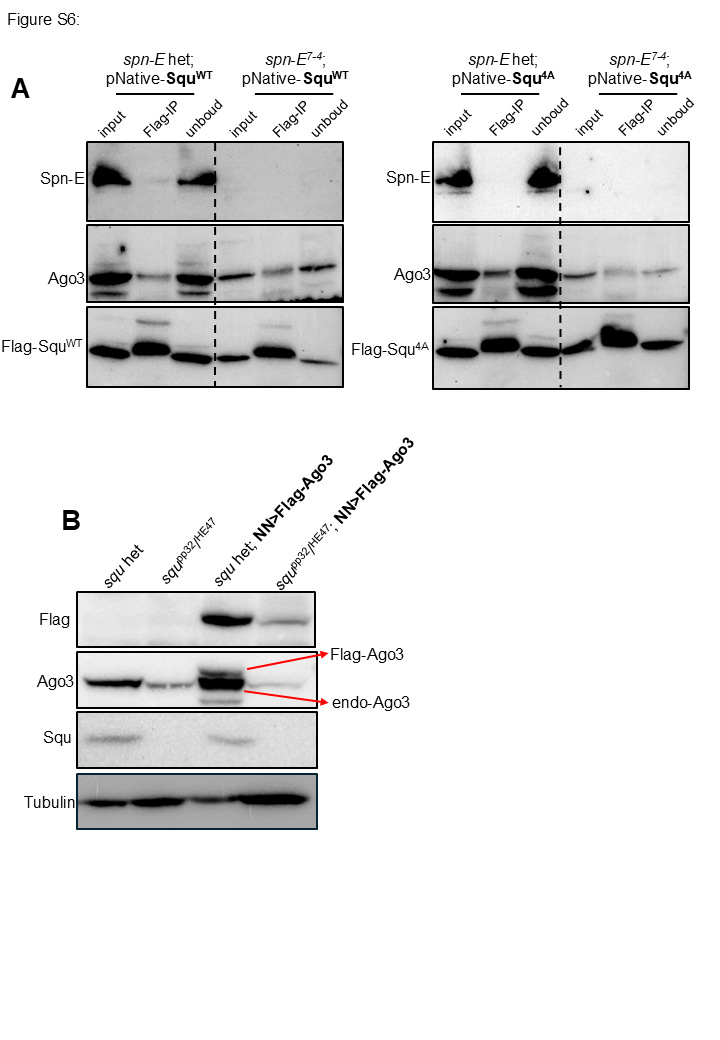

### Sup.Figure7

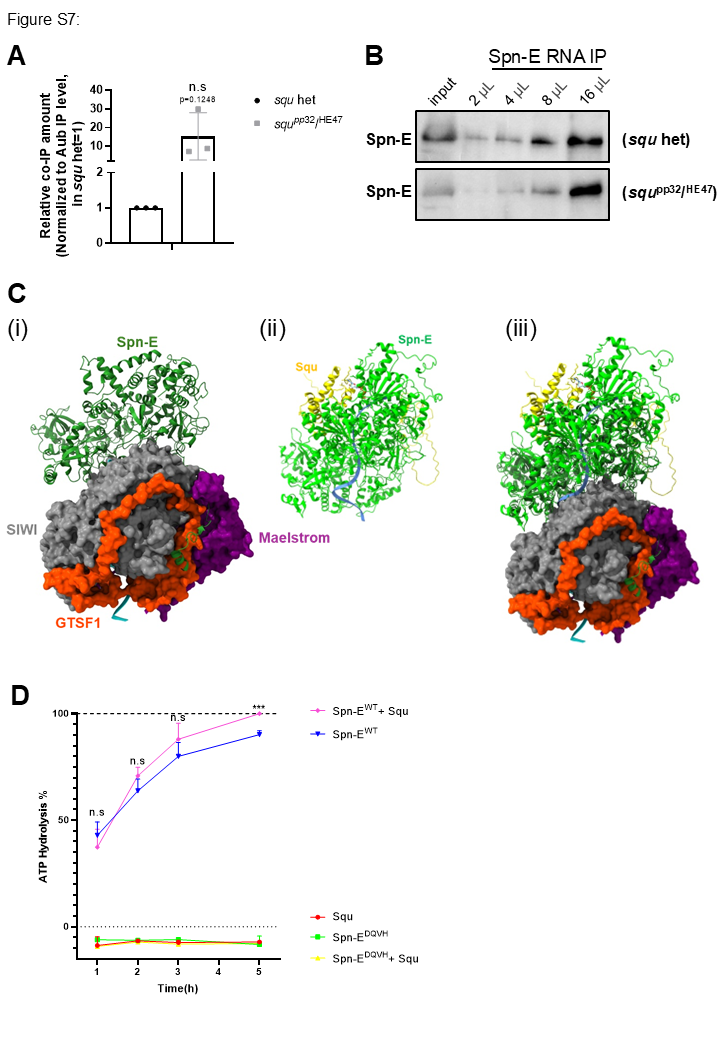
